# ER-dependent membrane repair of mycobacteria-induced vacuole damage

**DOI:** 10.1101/2023.04.17.537276

**Authors:** Aby Anand, Anna-Carina Mazur, Patricia Rosell-Arevalo, Rico Franzkoch, Leonhard Breitsprecher, Stevanus A. Listian, Sylvana V. Hüttel, Danica Müller, Deise G. Schäfer, Simone Vormittag, Hubert Hilbi, Markus Maniak, Maximiliano G. Gutierrez, Caroline Barisch

**Author notes:** **Corresponding author:** –.

## Abstract

Several intracellular pathogens, such as *Mycobacterium tuberculosis,* damage endomembranes to access the cytosol and subvert innate immune responses. The host counteracts endomembrane damage by recruiting repair machineries that retain the pathogen inside the vacuole.

Here, we show that the endoplasmic reticulum (ER)-Golgi protein oxysterol binding protein (OSBP) and its *Dictyostelium discoideum* homologue OSBP8 are recruited to the *Mycobacterium*-containing vacuole (MCV) after ESX-1-dependent membrane damage. Lack of OSBP8 causes a hyperaccumulation of phosphatidylinositol-4-phosphate (PI4P) on the MCV and decreased cell viability. OSBP8-depleted cells had reduced lysosomal and degradative capabilities of their vacuoles that favoured mycobacterial growth. In agreement with a function of OSBP8 in membrane repair, human macrophages infected with *M. tuberculosis* recruited OSBP in an ESX-1 dependent manner. These findings identified an ER-dependent repair mechanism for restoring MCVs in which OSBP8 functions to equilibrate PI4P levels on damaged membranes.

**Importance:** Tuberculosis still remains a global burden and is one of the top infectious diseases from a single pathogen. *Mycobacterium tuberculosis*, the causative agent, has perfected many ways to replicate and persist within its host. While mycobacteria induce vacuole damage to evade the toxic environment and eventually escape into the cytosol, the host recruits repair machineries to restore the MCV membrane. However, how lipids are delivered for membrane repair is poorly understood. Using advanced fluorescence imaging and volumetric correlative approaches, we demonstrate that this involves the recruitment of the ER-Golgi lipid transfer protein OSBP8 in the *D. discoideum*/ *M. marinum* system. Strikingly, depletion of OSBP8 affects lysosomal function accelerating mycobacterial growth. This indicates that an ER-dependent repair pathway constitutes a host defence mechanism against intracellular pathogens such as *M. tuberculosis*.

## Introduction

Cellular compartmentalization renders cells susceptible to membrane damage caused by pathogens, chemicals or mechanical stressors. Endolysosomal damage by vacuolar pathogens disrupts the proton gradient between the endolysosome and the cytosol and reduces the efficacy of first-line innate immune defences. Several pathogens including *Mycobacterium tuberculosis* have evolved sophisticated strategies to avoid phagosome maturation and to overcome the ion gradients (H^+^, Zn^2+^ or Cu^2+^), creating an optimal environment for their proliferation (1). Membrane damage inflicted by pathogenic mycobacteria depends on the pathogenicity locus region of difference (RD) 1 encoding the type VII secretion system ESX-1. This leads among others to the leakage of Zn^2+^ from the *Mycobacterium*-containing vacuole (MCV) thus preventing the bacteria from zinc poisoning (2).

Endosomal sorting complex required for transport (ESCRT)-dependent membrane repair plays a role during the infection of *Dictyostelium discoideum* with *M. marinum*, a pathogenic mycobacterium that primarily infects poikilotherms and is genetically closely related to the tuberculosis (TB) group (3). Importantly, the host response and course of infection by *M. tuberculosis* and *M. marinum* share a high level of similarity (4) including the molecular machinery for host lipid acquisition and turnover (5). In *M. marinum*-infected *D. discoideum*, the ESCRT machinery cooperates with autophagy to repair EsxA-mediated damage at the MCV (6). The evolutionarily conserved E3-ligase TrafE mobilises various ESCRT components to damaged lysosomes and the MCV (7). While the ESCRT components Tsg101, Chmp4/Vps32 and the AAA-ATPase Vps4 are recruited to small membrane ruptures, the autophagy machinery operates at places of extensive membrane damage (6). When ESCRT-dependent and autophagy pathways are disrupted, *M. marinum* escapes to the cytosol at very early infection stages, indicating that both mechanisms are needed to keep the bacteria inside the phagosome (6).

Two other repair pathways restore the integrity of broken lysosomal membranes in mammalian cells (8). Sphingomyelin (SM)-dependent repair operates at damaged lysosomes and ruptured vacuoles containing *Salmonella* Typhimurium (9) or *M. marinum* (10). Moreover, an endoplasmic reticulum (ER)-dependent membrane repair pathway has been described (11, 12). In this pathway, lysosomal damage results in the recruitment of PI4-kinase type 2-alpha (PI4K2A) generating high levels of phosphatidylinositol-4-phosphate (PI4P) (12). The accumulation of PI4P leads to the induction of ER-lysosome contacts and the mobilization of OSBP and several OSBP-related proteins (ORPs) that transfer cholesterol and phosphatidylserine (PS) from the ER to the ruptured lysosomes in exchange for PI4P (11, 12).

Proteomics and transcriptomics analyses indicate that ER-dependent membrane repair might also play a role during mycobacterial infection in human macrophages (13) and in *D. discoideum* (14, 15). Genes encoding the proteins involved in the establishment of membrane contact sites (MCS) with the ER or in lipid transfer are upregulated at infection stages, when major vacuolar damage occurs and the bacteria translocate to the cytosol. In this study, we investigated the role of OSBPs in ER-mediated membrane repair in the context of mycobacterial infection. We show that ESX-1- dependent membrane damage results in the mobilization of OSBP and its *D. discoideum* homologue OSBP8 to *M. tuberculosis*- and *M. marinum*-containing vacuoles, respectively. We demonstrate that OSBP8 is on ER-tubules in close contact with lysosomes and MCVs dependent on PI4P accumulation. OSBP8 depletion leads to cells that are less viable upon sterile damage. Upon infection, lack of OSBP8 causes a massive accumulation of PI4P on MCVs, impairs the functionality of this compartment and promotes mycobacterial replication. Altogether, our work reveals that OSBPs play an important role in equilibrating PI4P levels during ER-dependent repair to maintain the integrity of MCVs and contribute to the maintenance of the phagosomal innate immune defences against intracellular pathogens.

## Results

### Mycobacterial infection induces an ER-dependent repair gene expression signature

During lysosomal damage, cells stimulate a phosphatidylinositol (PI)-initiated signalling pathway for rapid lysosomal repair (12). This results in the recruitment of membrane tethers and lipid transfer proteins (LTPs) to ER-lysosome contact sites (11, 12). Analysis of RNA-sequencing data of *D. discoideum* during *M. marinum* infection (14, 15) revealed a possible role for ER-dependent repair: Genes encoding the homologues of the PI4P phosphatase Sac1 and several PI4Ks (*pi4k* and *pikD*) were upregulated at later infection stages when *M. marinum* inflicts major membrane damage (Fig. 1A). Additionally, the expression of many OSBPs is affected in complex manners during infection (Fig. 1B).

**FIG 1.**
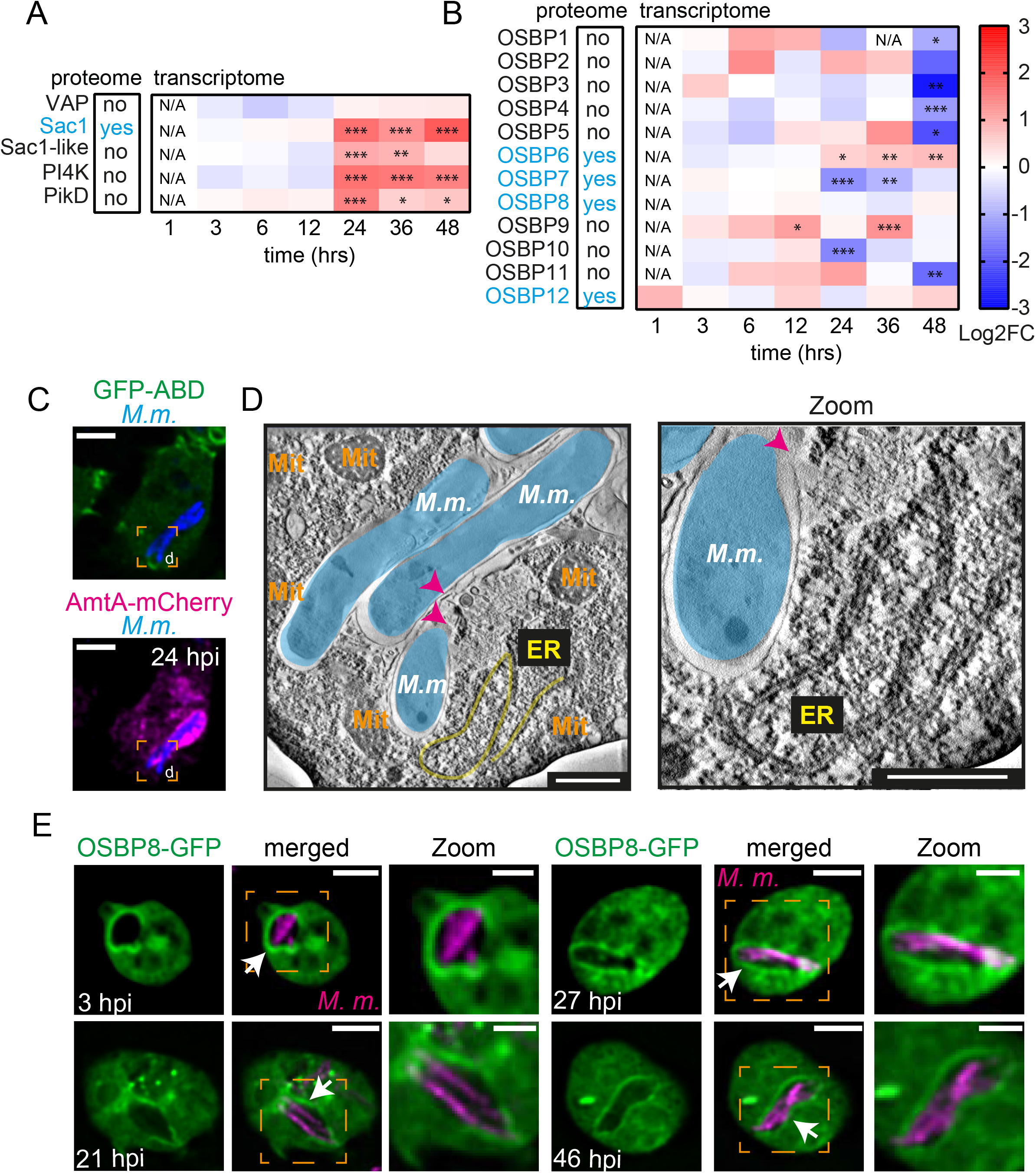
Evidence for ER-mediated repair during mycobacterial infection and mobilization of OSBP8. (A-B) Proteomics (left) and heatmaps (right) representing the transcriptional data derived from (14) and (15). Cells were infected with GFP-expressing *M. marinum* wt. Samples were collected at the indicated time points. Statistically significant differences in expression are marked with asterisks (*, P < 0.05; **, P < 0.01; ***, P < 0.001). Colours indicate the amplitude of expression (in logarithmic fold change (Log2FC)) in infected cells compared to mock-infected cells: from red (highest expression) to blue (lowest expression). (C-D) CLEM reveals ER-tubules close to ruptured MCVs. Cells expressing GFP-ABD and AmtA-mCherry were infected with eBFP-expressing *M. marinum*. At 24 hpi, cells on sapphire discs were imaged by SD microscopy in the presence of low concentrations of GA before high-pressure freezing. Left: deconvolved SD images, scale bars, 5 µm; right: representative EM micrographs, scale bars, 500 nm. Magenta arrow heads point to the ruptured MCV membrane. Mitochondria (Mit) were pseudo-coloured in orange, *M. marinum* (*M.m*.) in cyan and ER-tubules in yellow. (E) OSBP8-GFP is recruited to intracellular mycobacteria. Cells overexpressing OSBP8-GFP were infected with mCherry-expressing *M. marinum*. At the indicated time points, cells were imaged live by spinning disc (SD) microscopy. Arrows point to OSBP8-GFP^+^ mycobacteria. Scale bars, 5 µm; Zoom, 2 µm. Images in (C and E) were deconvolved.

In *D. discoideum*, *M. marinum* resides in a compartment with partially lysosomal and post-lysosomal characteristics that is exposed to damage starting from early infection stages (6, 16). We investigated whether mycobacterial infection leads to the formation of ER-MCV contacts. Indeed, when cells expressing the ER-marker Calnexin-mCherry were infected with *M. marinum* and stained for the MCV-marker p80 (16), we observed Calnexin^+^ ER-tubules in the close vicinity of the MCV (Fig. S1A). This is consistent with previous findings showing *M. tuberculosis* infection of dendritic cells, in which approximately 50% of the MCVs were Calnexin^+^ (17). To gain a better understanding of the morphology of these sites, cells expressing GFP-actin-binding-domain-(ABD) as well as the endosomal and MCV marker AmtA-mCherry (18) were infected and analysed by correlative light and electron microscopy (CLEM). The overexpression of GFP-ABD leads to a significant improvement of cell adhesion and was necessary to re-locate the cells after sample preparation for EM. By correlating the images of vacuolar bacteria obtained by live cell imaging (AmtA^+^) (Fig. 1C) with the corresponding EM micrographs, ER-tubules were seen close to ruptured MCVs (Fig. 1D).

In summary, we discovered a unique transcriptomic signature that, together with the observation of ER-tubules in the proximity of the MCV, supports the hypothesis that mycobacterial infection triggers ER-dependent membrane repair.

### OSBP8-GFP is mobilized by intracellular mycobacteria

LTPs from the OSBP/ORP family are mobilized during ER-dependent lysosomal repair to provide lipids such as PS or cholesterol. Members of this protein family counter-transport these lipids in exchange for PI4P at ER-lysosome contacts. Sequence comparison of the twelve *D. discoideum* OSBPs with human and yeast homologues revealed that family members consist primarily of the lipid-binding OSBP-related domain (ORD) (Fig. S1B) (19). By analysing recent proteomics data from infected *D. discoideum* (14), we found four out of the twelve *D. discoideum* homologues enriched on candidates for OSBP-mediated ER-dependent repair during infection. The fact that OSBP8 is the closest homologue to mammalian family members (19) and is the only *D. discoideum* OSBP with a fully conserved EQVSHHPP lipid-binding motif (Fig. S1C) prompted us to investigate its localization during mycobacterial infection and to use OSBP7, which is more distantly related, as a control.

To study the subcellular localization of OSBP8, we overexpressed it tagged with GFP at either end (Fig. S2A-F). OSBP8-GFP was partly cytosolic and co-localized with Calnexin-mCherry at the perinuclear ER as well as with ZntC-mCherry, a zinc transporter that is located at the Golgi and/or recycling endosomes (20) (Fig. S2A-B). Interestingly, the membrane localization of OSBP8 was abolished in cells overexpressing GFP-OSBP8, indicating that the N-terminus is important for membrane targeting (Fig. S2C-D).

Strikingly, when the localization of OSBP8 was monitored during infection with *M. marinum*, OSBP8- GFP re-localized to MCVs starting from early stages (Fig. 1C). In contrast, OSBP8-GFP did not localize on bead-containing phagosomes (BCPs) (Fig. S2E-F), indicating a specific response to vacuoles containing mycobacteria. OSBP7 localized in the cytosol and nucleus in non-infected cells and did not re-localise during infection (Fig. S2G) suggesting that some OSBPs are specifically mobilized in response to infection.

Overall, OSBP8 is specifically recruited to MCVs starting from early infection stages, which correlates with the occurrence of MCV-damage in *D. discoideum* (6).

### OSBP8-GFP is located on ER-tubules in the vicinity of damaged MCVs

To test if OSBP8-GFP was recruited to MCVs or cytosolic mycobacteria, we infected cells expressing OSBP8-GFP and AmtA-mCherry and performed lattice light sheet microscopy (LLSM) (Fig. 2A-C). OSBP8-GFP did not colocalize with the MCV membrane (AmtA^+^), but it was recruited to its immediate vicinity (Fig. 2A-B; see also Movie S1). This finding was further corroborated by a 3D analysis that demonstrates that some MCVs were fully enclosed by OSBP8-GFP^+^ structures (Fig. 2C; see also Movie S2). To visualize these potential ER-MCV contacts, expansion microscopy (ExM) of cells expressing OSBP8-GFP and Calnexin-mCherry was carried out. In line with the previous results, Calnexin-mCherry^+^ and OSBP8-GFP^+^ ER-tubules were observed in the vicinity of the MCV (Fig. 3D). Additionally, we performed CLEM (Fig. S3A-B) and 3D-CLEM (Fig. 2E-G, Fig. S3C) to acquire a deeper insight of the morphology of these micro-compartments. The CLEM analysis confirmed by live imaging that OSBP8-mCherry is recruited to the MCV. Strikingly, in the corresponding EM images OSBP8-mCherry coincides with ER-tubules that were in the vicinity of seemingly damaged MCVs (Fig. S3A-B). For volumetric image analysis, the infected cells were subjected after live cell imaging to serial block face-scanning electron microscopy (SBF-SEM). In agreement with our previous observations, electron micrographs and the 3D rendering clearly showed that the MCV is surrounded by OSBP8^+^-ER-tubules (Fig. 2E-G, Fig. S3C; see also Movie mycobacterial infection.

**FIG 2.**
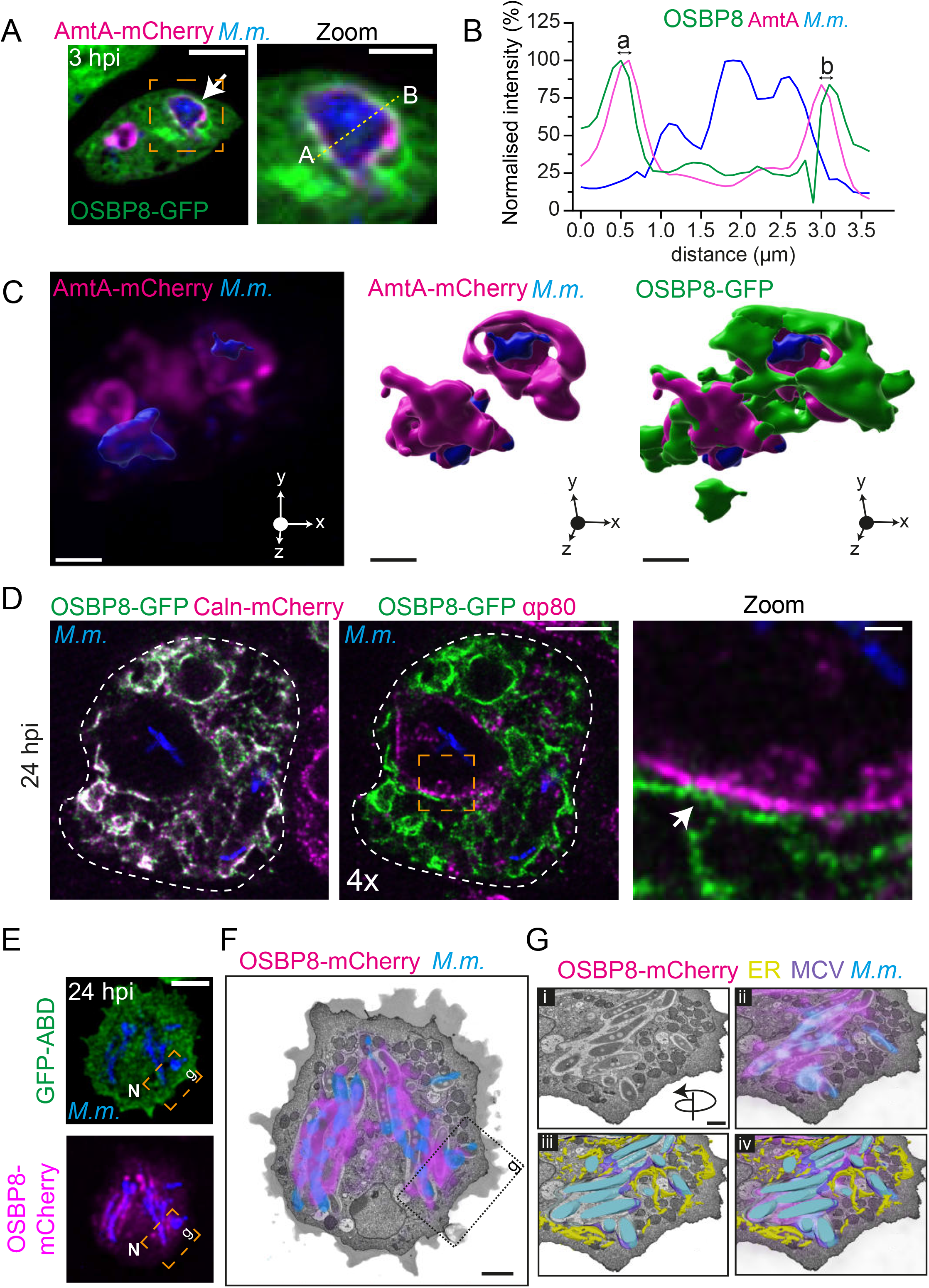
OSBP8-GFP is located on ER-tubules in the vicinity of the MCV. (A) OSBP8-GFP localizes adjacent to the MCV membrane. (B) Intensity profile of the line plotted through the MCV shown in the zoom of A. (C) 3D-model of the cell shown in (A) illustrating OSBP8-GFP^+^ membranes capping the MCV (AmtA^+^). Cells dually expressing OSBP8-GFP/AmtA-mCherry were infected with eBFP-expressing *M. marinum* and imaged live at 3 hpi by LLSM. Arrow points to OSBP8-GFP^+^ membranes close to the MCV. Scale bars in A, 5 µm; Zoom, 2 µm; in C, 2 µm. (D) OSBP8-GFP is located on ER-tubules in the proximity of the MCV. Cells dually expressing OSBP8-GFP/Calnexin-mCherry were infected with eBFP-expressing *M. marinum*, fixed at 24 hpi and stained with antibodies against p80, GFP and mCherry before 4x expansion. Arrow points to an OSBP8-GFP^+^ ER-tubule close to localized on ER-tubules close to the MCV (arrow). Cells expressing OSBP8-mCherry/GFP-ABD were infected with eBFP-expressing *M. marinum*. At 24 hpi, cells were imaged by SD microscopy (E) and prepared for SBF-SEM (F-G). (F) EM micrograph showing the cell with the correlated OSBP8-mCherry and eBFP-*M. marinum* signal. Please see Fig. S3C for more information. (G) Closeup of the position indicated in F showing ER-tubules close to the MCV. (i-iv) correlation of the (i) EM micrograph (ii) with OSBP8-mCherry (magenta) and mycobacteria (blue). (iii-iv) segmentation of the ER (yellow), MCV (violet) and mycobacteria (cyan). Scale bars, 5 µm (E); 2 µm (F) and 1 µm (G). SD images were deconvolved. N: nucleus.

**FIG 3.**
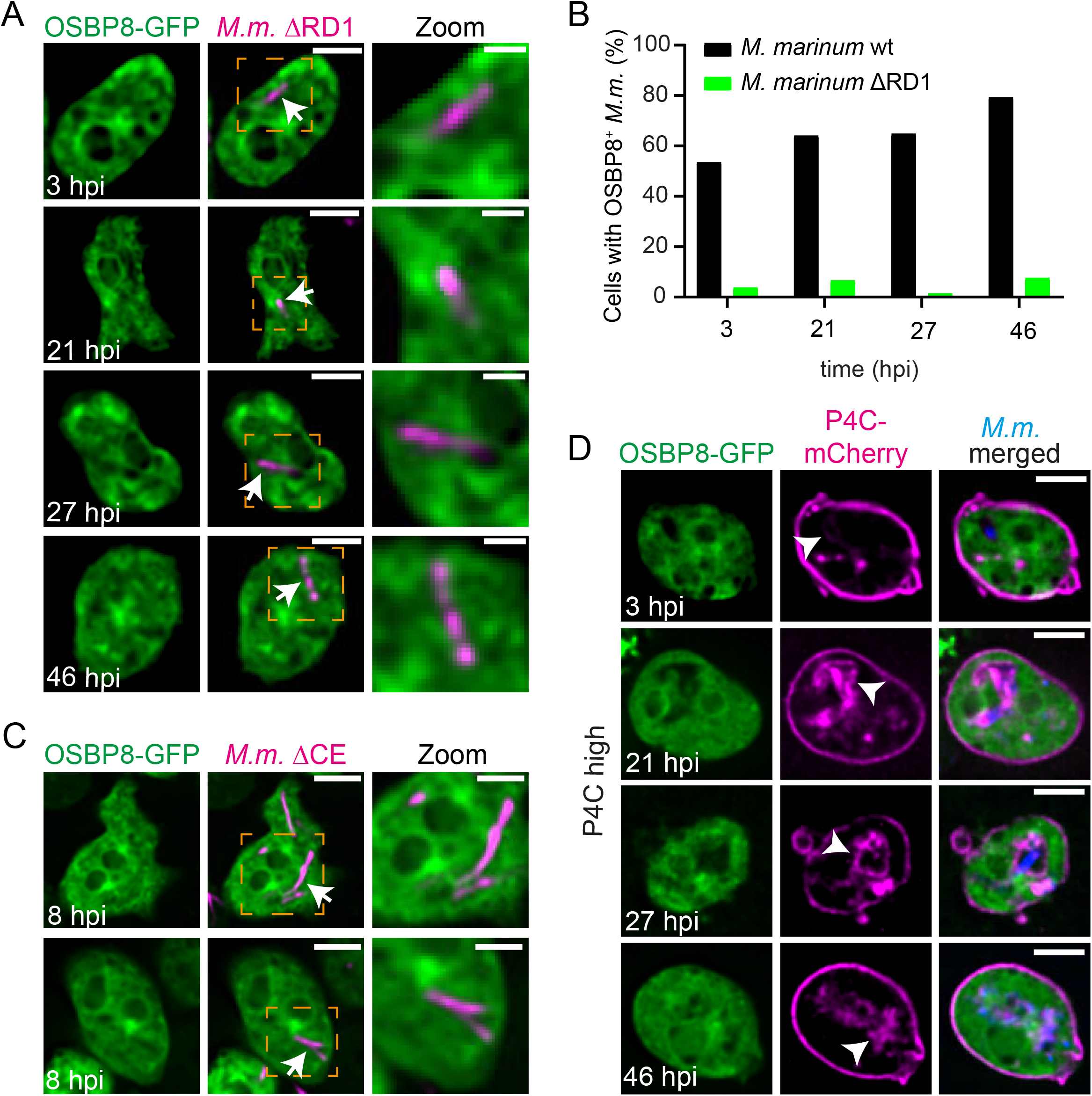
OSBP8-GFP mobilization during infection is dependent on ESX-1/EsxA and PI4P. (A) OSBP8- GFP is not recruited to intracellular *M. marinum* ΔRD1 mutant. (B) Quantification of A. Data represent two independent experiments (OSBP8-GFP 3, 21, 27, 46 hpi N=2, 23≤n≤274). (C) OSBP8-GFP is not recruited to ΔCE mutant. (D) OSBP8-GFP is not mobilized during infection in cells highly expressing P4C-mCherry. Cells overexpressing OSBP8-GFP or co-expressing P4C-mCherry were infected with mCherry-or eBFP-expressing *M. marinum* wt, ΔRD1 or ΔCE. At the indicated time points samples were taken for SD microscopy. Arrows point to OSBP8-GFP^-^ intracellular mycobacteria. Arrow heads indicate PI4P^+^ MCV. Scale bars, 5 µm; Zoom, 2.5 µm. Images were deconvolved. *M.m.: M. marinum*.

Next, we tested if the mobilization of OSBP8 to the MCV is damage-dependent and performed infections with a mutant of *M. marinum* lacking ESX-1 (ΔRD1) (6). Remarkably, the localization of OSBP8-GFP in the vicinity of the MCV was abolished in cells infected with the ΔRD1 mutant (Fig. 3A-B). A similar observation was made with the ΔCE mutant that has a functional ESX-1 system but lacks EsxA together with its chaperone EsxB (Fig. 3C). Since overexpression can lead to artefacts and to the induction of MCS, we generated a chromosomally-tagged GFP-fusion of OSBP8 (OSBP8::GFP) in which OSBP8 is under the control of its endogenous promoter and expressed 20 times less than in OSBP8-GFP overexpressing cells (Fig. S4A-B). OSBP8::GFP shows a similar distribution and is also recruited to the MCV in an ESX-1-dependent manner (Fig. S4C-D). We concluded that ESX-1/EsxA-mediated membrane damage triggers the formation of ER-MCV MCS and the recruitment of OSBP8-GFP to these sites.

### The recruitment of OSBP8-GFP to damaged lysosomes and the MCV is dependent on PI4P

We analysed the distribution of OSBP8-GFP upon treatment with lysosome disrupting agent Leu-Leu-O-Me (LLOMe) (21) to test whether OSBP8 is recruited as a general response to lysosomal damage. As observed previously, LLOMe induced the formation of ESCRT-III-(GFP-Vps32^+^) structures at the periphery of lysosomes labelled with fluorescent dextran (Fig. S5A) (6). In mammalian cells, ER-dependent lysosome repair is initiated by the recruitment of PI4K2A leading to an accumulation of PI4P at the damage site and the recruitment of ORPs/OSBPs (11, 12). Also in *D. discoideum,* PI4P, visualized with the PI4P-binding domain (P4C) of the *Legionella* effector SidC (22-24), was rapidly observed on ruptured lysosomes. The kinetics of P4C-GFP associated with damaged lysosomes was slightly delayed compared to GFP-Vps32 (Fig. S5A). Also OSBP8- GFP was mobilized upon sterile damage (Fig. S5A), however, as observed for OSBP in HeLa (11) and U2OS cells (12), the recruitment happened relatively late (20-40 min after LLOMe treatment) and was observed less frequently. In contrast, OSBP7-GFP remained cytosolic and in the nucleus upon LLOMe treatment (Fig. S5A). This implies that OSBP8-GFP^+^ ER-lysosome contacts were a response to lysosomal damage and that this pathway might be activated after SM-and ESCRT-dependent repair. Intriguingly, the mobilization of OSBP8-GFP was totally abolished in cells highly expressing P4C-mCherry (Fig. S5B), indicating that P4C binds to PI4P with such a high affinity that it displaces OSBP8-GFP. In cells expressing P4C-mCherry at low levels, OSBP8-GFP became visible at the periphery of dextran-labelled endosomes upon LLOMe treatment (Fig. S5C). A similar observation was made during infection: Here, P4C-mCherry accumulated at the MCV starting from early infection stages. In cells highly expressing P4C-mCherry, OSBP8-GFP was not recruited to the MCV (Fig. 3D), indicating that P4C-mCherry competes with OSBP8-GFP for PI4P. However, when P4C-mCherry was expressed at a low level, OSBP8-GFP co-localized with *M. marinum* (Fig. S5D). Altogether, these data indicate that OSBP8-GFP is recruited by PI4P on damaged lysosomes and on ruptured MCVs, respectively.

### OSBP8 prevents accumulation of PI4P on damaged MCVs and restricts mycobacterial growth

OSBP8 localizes to the perinuclear ER and the juxtanuclear region that is characteristic for the Golgi apparatus in *D. discoideum* and might be a homologue of mammalian OSBP that is recruited to ER-Golgi contacts to shuttle PI4P and cholesterol between the two organelles (25). In ER-dependent membrane repair, OSBP was reported to balance out PI4P levels on ruptured lysosomes (11). To investigate if OSBP8 is involved in PI4P transport, we generated knockouts (KOs), in which the corresponding gene (*osbH*) is disrupted by a BS^r^ cassette. Similar to OSBP in mammalian cells, deletion of OSBP8 caused a re-distribution of the PI4P probe away from the PM to internal structures reminiscent of the Golgi apparatus (Fig. S6A-B). Upon LLOMe treatment, P4C-GFP^+^ lysosomes were detected for up to 110 min in cells lacking OSBP8, whereas the P4C-GFP signal dissociated from lysosomes of wild type (wt) cells considerably earlier (Fig. S6C). Additionally, as observed for OSBP (11), OSBP8 was essential for cell viability following LLOMe treatment, however, the effect was less pronounced compared to cells lacking Tsg101 (Fig. S6D-E). Taken together, our data provide strong evidence that OSBP8 is involved in PI4P-removal from ruptured lysosomes.

During infection, we observed a hyperaccumulation of P4C-GFP on the MCVs of the *osbH* KO (Fig. 4A-B). Previous data by us and others indicate that sterols accumulate in the MCV of *M. marinum* (18) and *M. bovis* BCG (26). To test if OSBP8 depletion interferes with sterol transport, we performed filipin staining. A statistically significant lower filipin intensity was observed on MCVs in *osbH* KO at later infection stages (Fig. 4C-D), suggesting that sterols might indeed shuttled by OSBP8. Since the difference was small, sterols might be additionally transferred to the MCV by other mechanisms. PI4P accumulation on lysosomes of OSBP-depleted cells was hypothesized to induce increased and prolonged ER-endosome contact sites and might impact on lysosomal function (11). During infection approximately half of the bacteria were positive for the peripheral subunit VatB of the H^+^-ATPase and the MCVs are labelled by LysoSensor and DQ-BSA at early infection stages (27). Depletion of OSBP8 lead to a decrease of LysoSensor^+^ bacteria and to MCVs that are less acidic (Fig. 4E-G). Although the percentage of DQ-BSA^+^ bacteria was equivalent to wt, MCVs in *osbH* KOs were less degradative (Fig. 4H-J), indicating that the increased accumulation of PI4P impaired the lysosomal and proteolytic capabilities of the MCV. This finding further supported by the fact that intracellular *M. marinum* growth was increased in two independent *osbH* KOs (Fig. 4K). Conversely, overexpression of OSBP8-GFP lead to reduced intracellular growth (Fig. 4l). Importantly, deletion of OSBP8 did not impact vacuolar escape (Fig. S6F-G) suggesting that the MCV environment is responsible for the growth advantage of the bacteria.

**FIG 4.**
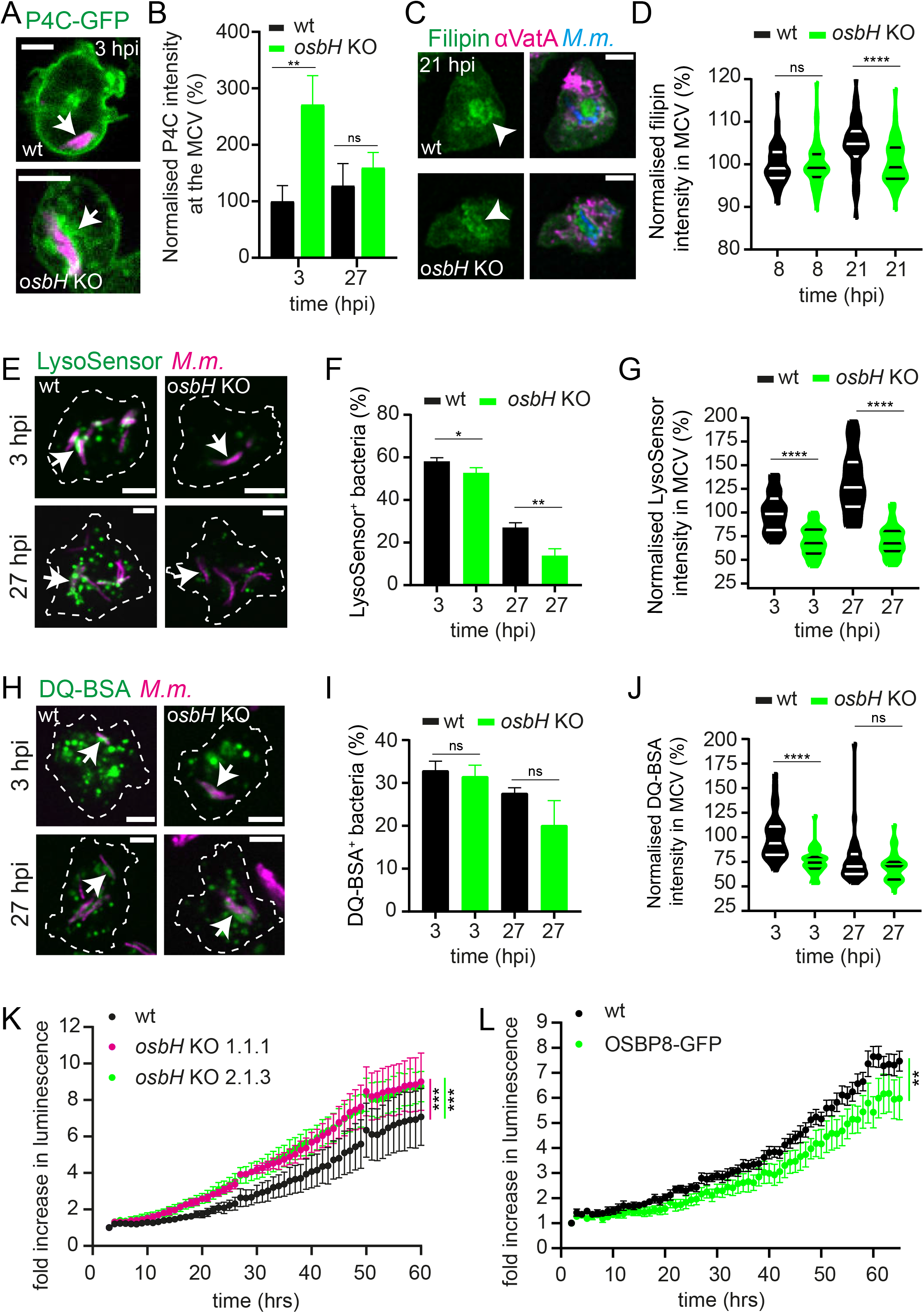
OSBP8 prevents PI4P accumulation on MCVs while maintaining their lysosomal and degradative properties. (A) P4C-GFP hyperaccumulates on MCVs of the *osbH* KO. (B) Quantification of A. Data are representative for one of two independent experiments (P4C-GFP 3, 27 hpi N=2, 14≤n≤24). (C) Sterol distribution in wt vs. *osbH* KO cells infected with *M. marinum* wt. (D) Quantification of C. Plots show the mean and standard deviation of three independent experiments (Filipin 8, 21 hpi N=3, 30≤n≤70). Statistical differences were calculated with an unpaired t test (*p < 0.05; **** p < 0.0001). (E) Lysosomal properties of MCVs in wt vs. *osbH* KO cells. (F-G) Quantifications of E. Plots show the mean and standard deviation of three independent experiments (LysoSensor green 3, 27 hpi N=3, 290≤n≤450). Statistical differences were calculated with an unpaired t test (*p < 0.05; **p < 0.01; **** p < 0.0001). (H) Proteolytic activity of MCVs in wt vs. *osbH* KO cells. (I-J) Quantification of H. Plots show the mean and standard deviation of three independent experiments (DQ-BSA green 3, 27 hpi N=3, 140≤n≤240). Statistical differences were calculated with an unpaired t test (**** p < 0.0001). *D. discoideum* wt and *osbH* KO (expressing P4C-GFP (a)) were infected with mCherry-or eBFP-expressing *M. marinum* wt. At the indicated time points samples were taken for SD microscopy. For filipin staining, infected cells were fixed and stained for VatA. Arrows point to PI4P^+^ or LysoSensor^+^ or DQ-BSA^+^ MCV and arrow heads indicate sterol accumulation in the MCV. Scale bars, 5 µm. Images in C were deconvolved. *M.m.: M. marinum.* (K-L) Mycobacterial growth is altered in cells lacking OSBP8 or cells overexpressing OSBP8-GFP. *D. discoideum* wt, two independent *osbH* KOs or OSBP8-GFP overexpressing cells were infected with *M. marinum* wt expressing bacterial luciferase. Luminescence was recorded every hour with a microplate reader. Shown is the fold increase in luminescence over time. Symbols and error bars indicate the mean and SEM of three independent experiments. Statistical differences of pairwise comparisons were calculated with a Fisher LSD post hoc test after two-way ANOVA (**, P < 0.01; ***, P < 0.001).

Thus, OSBP8-mediated removal of PI4P from the MCV during ER-dependent membrane repair is necessary to preserve the membrane integrity and the lysosomal functionality of this compartment.

### *M. tuberculosis* recruits OSBP in an ESX-1-dependent manner in human macrophages

Next, we sought to validate our findings in induced pluripotent stem cell (iPSC)-derived macrophages (iPSDMs) infected with *M. tuberculosis* (Fig. 5). According to RNA-sequencing, key genes of this repair pathway were significantly upregulated in an ESX-1-dependent manner (Fig 5A-B). We observed a higher expression level of *ORP5*, *ORP9* and *PI4K3B,* i.e. another PI-kinase that also localises to lysosomes (28). This signature was significant at 48 hpi and most of the genes were not upregulated in cells infected with the *M. tuberculosis* ΔRD1 mutant. OSBP transfers cholesterol to damaged lysosomes to preserve membrane stability and PI4P in the opposite direction to ensure the establishment of functional contact sites (11). Strikingly, during infection of iPSDMs, endogenous OSBP re-localized to *M. tuberculosis* wt (Fig. 5C). Notably, this recruitment is ESX-1 dependent as OSBP was less efficiently recruited in cells infected with *M. tuberculosis* ΔRD1 mutant (Fig. 5D). Collectively, our data highlight the evolutionary conservation of an ER-dependent membrane repair mechanism from simple eukaryotes such as *D. discoideum* to human cells.

**FIG 5.**
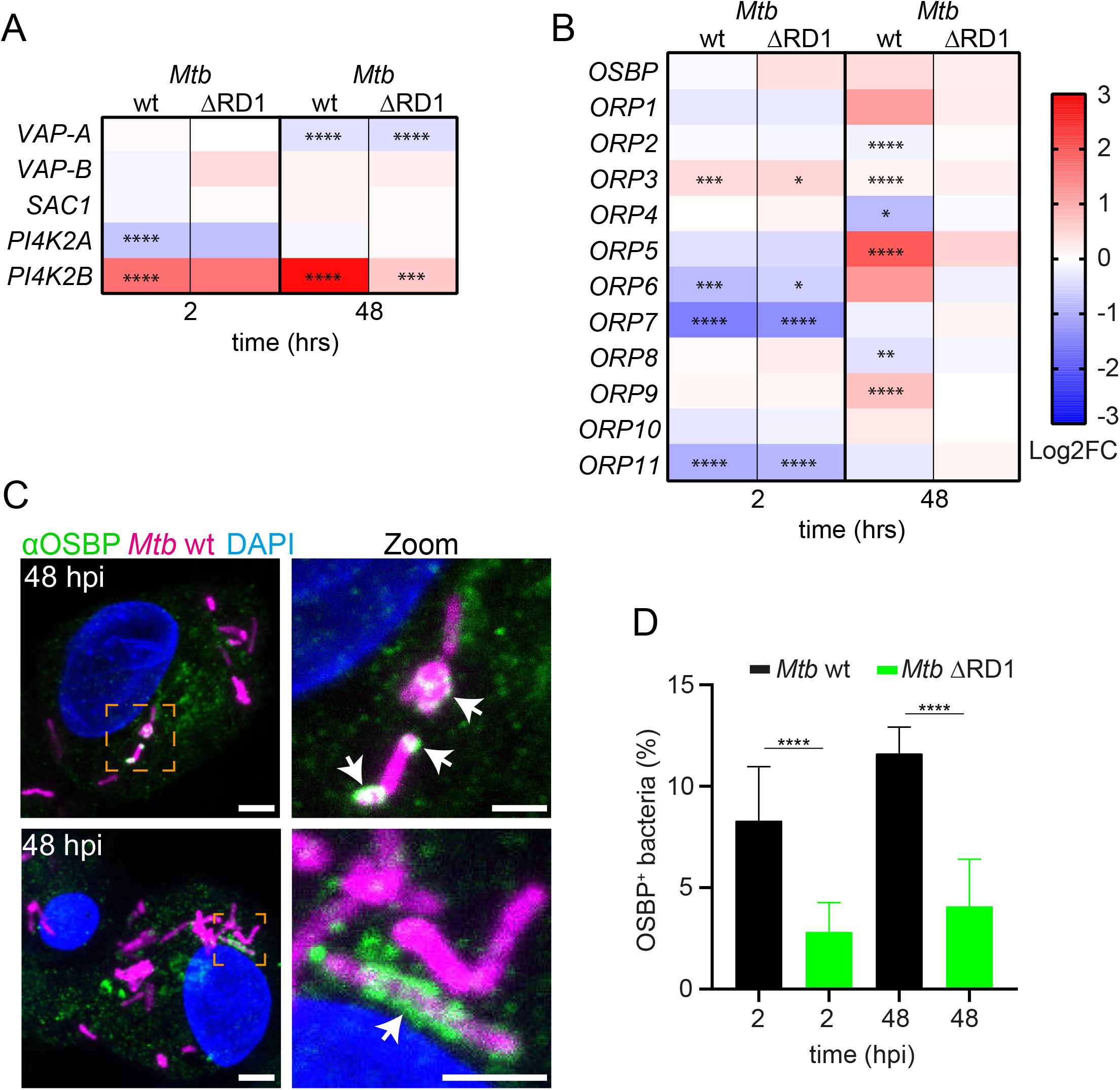
ER-mediated repair plays a role during *M. tuberculosis* infection. (A-B) Heatmaps of differentially expressed genes (Log_2_ Fold-change values) encoding proteins involved in MCS formation from RNA-sequencing analysis of human iPSDMs infected with either *M. tuberculosis* wt or ΔRD1 (13). Samples were collected at the indicated time points. Statistically significant differences in expression are marked with asterisk (*, P < 0.05; **, P < 0.01; ***, P < 0.001, ****, P < 0.0001). Colours indicate the amplitude of expression (Log2FC) in infected cells compared to mock-infected cells: from red (highest expression) to blue (lowest expression). Data was retrieved from (13). (C) In human iPSDMs OSBP is recruited to *M. tuberculosis* wt but not to the ΔRD1 mutant. iPSDMs were infected with E2-Crimson-expressing bacteria. At 2 and 48 hpi cells were fixed and stained against OSBP. Shown are two representative images from 48 hpi. Z-stacks: 20, 0.3 μm. Scale bars, 5 μm; Zoom; 2 μm. (D) Quantification of C. Plots show the mean and standard deviation of three independent experiments (OSBP 2, 48 hpi N=3, 800 ≤n≤1200). Statistical differences were calculated with an unpaired t-test (**** P < 0.0001). *Mtb*: *M. tuberculosis*.

## Discussion

Using transcriptomics and proteomics data of infected cells as well as advanced imaging approaches, we provide evidence that ER-dependent repair is involved in mycobacterial infection. The main features of this membrane repair pathway at MCVs are shown in Fig. 6. Since various genes encoding for PI4Ks are upregulated at later infection stages (Fig. 1A), we hypothesize that cumulative damage at the MCV leads to the recruitment of PI4K. This is consistent with the fact that PI4P accumulated at this compartment (Fig. 3D). The presence of PI4P is essential for (i) the formation of MCS via the interaction with PI4P-binding, tethering proteins that might interact with the anchor VAP at the ER and (ii) for the recruitment of LTPs belonging to the OSBP/ORP-family. We suggest that lipid transport is fuelled by a PI4P gradient that is maintained by the PI4P hydrolase Sac1 on the ER. Conversely, an upregulation of Sac1 during infection was observed in *D. discoideum* (Fig. 1A). We propose that the ER-dependent pathway plays a role in providing lipids for other membrane repair mechanisms, including SM-and ESCRT-dependent repair (8). In line with that we also observed an upregulation of *ORP5* and *ORP9* during *M. tuberculosis* infection. The corresponding proteins might transfer PS to the MCV (12, 29). These lipids might be essential for the generation of intraluminal vesicles, which ultimately facilitate the removal of the damage site. To better understand the potential crosstalk between SM-, ESCRT-, and ER-dependent repair mechanisms during infection further work is necessary.

**FIG 6.**
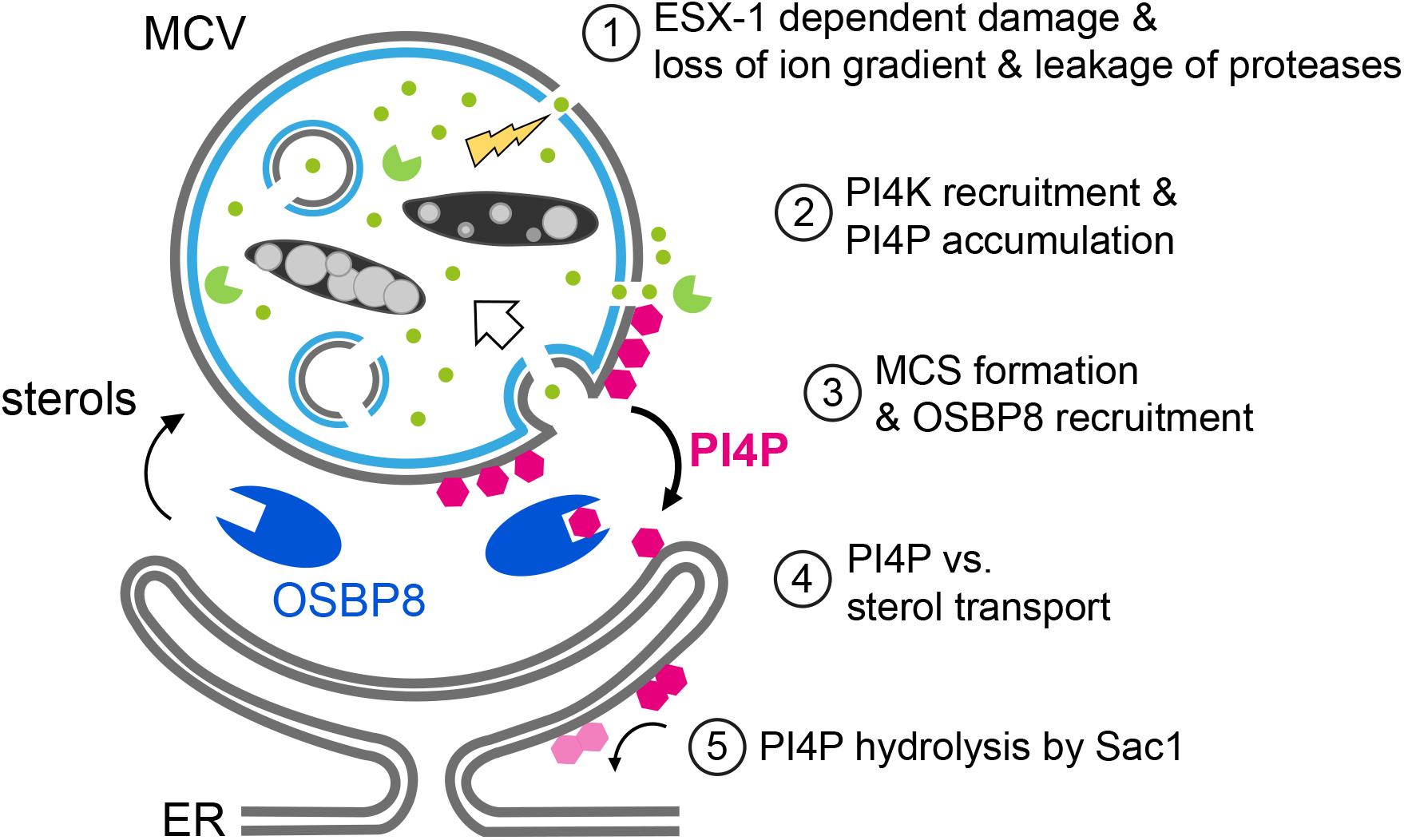
Schematic outline of ER-dependent repair during mycobacterial infection. 1.) ESX1- dependent vacuolar damage (yellow flash) leads to a loss of ion gradients (green spots) and the release of proteases (green packmen). 2.) PI4K are recruited to generate PI4P (pink polygons) at the MCV. 3.) This leads to the establishment of ER-MCV-MCS and the mobilization of OSBP8 (blue) in *M. marinum*-infected *D. discoideum*. 4.) OSBP8 transports sterols from the ER to the MCV and PI4P in the opposite direction. 5.) The transport is fuelled by the lipid phosphatase Sac1 that hydrolyses PI4P generating PI (light pink).

Besides transferring sterols to the MCV, OSBP8 and mammalian OSBP have a crucial role in equilibrating PI4P levels to ensure the formation of functional ER-MCV MCS. Strikingly, both proteins, were recruited to MCVs. Mycobacteria lacking ESX-1 failed to mobilize these proteins (Fig. 3A-C, Fig. 5C-D), indicating that membrane damage is a prerequisite for their recruitment. Using advanced imaging approaches such as LLSM, CLEM and 3D-CLEM, we discovered that OSBP8 is on ER-tubules that form MCS with the MCV (Fig. 2, Fig. S3). OSBP8 depletion resulted in the hyperaccumulation of PI4P on MCVs (Fig. 4A-B) and intracellular growth of *M. marinum* was accelerated (Fig. 4K). The observed growth advantage is probably due to the impaired lysosomal function and degradative properties of the MCV in the absence of OSBP8 (Fig. 4E-J). Depletion of OSBP8 did not fully inhibit the accumulation of sterols inside the MCV (Fig. 4C-D), thus, sterol transport might be mediated either by vesicular transport or other sterol transporters.

How is OSBP8 recruited to ER-MCV MCS? Members of the OSBP/ORP family are typically targeted to the ER by binding to VAP through their two phenylalanines-in-an-acidic tract (FFAT)-motif. Sequence analysis revealed that all *D. discoideum* OSBPs are short and contain neither a FFAT-motif, nor pleckstrin homology- (PH-) or transmembrane domains but consist primarily of the ORD (Fig. S1B). OSBP8 has a short amphipathic lipid packing sensor (ALPS)-like motif (30) flanked by an unstructured N-terminus (Fig. S6H). Intriguingly, the presence of an N-terminal GFP prevented membrane targeting of OSBP8 (Fig. S2C-D) suggesting that the ALPS-like motif may be involved in PI4P-binding (Fig. 3D, Fig. S2C-D).

In summary, *D. discoideum* and macrophages restrict pathogenic mycobacteria such as *M. tuberculosis* and *M. marinum* by restoring the MCV membrane with the help of ER-dependent membrane repair. We conclude that PI4P levels at the MCV need to be tightly regulated to allow the correct establishment of ER-MCV MCS to provide adequate levels of lipids to preserve membrane integrity. This in turn is necessary to maintain ion gradients and fundamental innate immune functions of these compartments. Our findings pave the way for an in-depth mechanistic analysis of the role of ER-dependent repair for the formation and stability of pathogen vacuoles.

## Materials

### D. discoideum plasmids, strains and cell culture

All the *D. discoideum* material is listed in Table S1. *D. discoideum* wt (AX2) was grown axenically at 22°C in Hl5c medium (Formedium) containing 100 U/mL penicillin and 100 µg/mL streptomycin. To generate *osbH* KOs, *osbH* was amplified with the primers #293 (5’ CGG AAT TCA AAA TGT TTT CAG GAG CAT TG) and #294 (5’ CGG AAT TCT TAA TTT GAA GCT GCT GC) from genomic DNA of AX2 digested with EcoRI and ligated into the same site of pGEM-T-Easy (Promega) to yield plasmid #625. From this plasmid a central 0.3 kbp fragment was eliminated by MfeI and the ends were blunted by T4 DNA polymerase. Thereafter the blasticidin S-resistance cassette flanked by SmaI sites from plasmid pLPBLP (31) was inserted, resulting in plasmid #629. Digestion with EcoRI produced an *osbH* gene interrupted by the BS^r^-cassette that was used for electroporation. The *D. discoideum* clones were screened and verified by PCR with #353 (5’ CAAT ACC AAT AGA TTT TAT ATC ATT AC) that bound genomic DNA just upstream the construct used for targeting and primers #57 (5’ CGC TAC TTC TAC TAA TTC TAG A) complementary to the 5’ end of the resistance cassette. Because this primer combination did not yield a product for the wildtype, further verifications involved primers #353 in combination with #358 (5’ CCT CTG ATG AGT TAC CAT AG) in the 3’ homologous sequence, as well as #357 (5’ GCC TCA AAA CAA GAT AGC G) binding in the 5’ region of the targeting construct together with #356 (5’ CAG CGG AAA TTG AAT GAA TAA ATT) complementary to a sequence downstream of the region used for homologous recombination. The OSBP8::GFP knockin cell line was generated using a previously described strategy (32) with the aim to insert the GFP-tag and the blasticidin cassette after the endogenous gene by homologues recombination. To this end, two recombination arms consisting of the last ∼ 500 bp of *osbH* (left arm (RA)) and of ∼ 500 bp downstream of *osbH* (right arm (RA) were amplified by PCR using the LA primers (oMIB56: 5’CGA GAT CTG GTT GGT TAG GTG CCG GTC G and oMIB57: 5’GGA CTA GTA TTT GAA GCT GCT GCT TTA ACT CTT TCT TCT C) as well as the RA primers (oMIB101: 5’CGG TCG ACT AAA AAC AAT AAT AAT TAT ATA TTT TAA TCG TAA ACA ATT TAT TCA TTC AAT TTC C and oMIB102: 5’GCG AGC TCG GAA ATC TTG TTG GAG G) and cloned into the plasmid pPI183 (32) using the restriction sites BglII, BcuI (LA) and SalI and SacI (RA). The resulting plasmid pMIB173 was used for electroporation after linearization with PvuII. The *D. discoideum* clones were screened and verified by PCR with the primers oMIB56 (5’CGA GAT CTG GTT GGT TAG GTG CCG GTC G) and oMIB57 (5’ GGA CTA GTA TTT GAA GCT GCT GCT TTA ACT CTT TCT TCT C) complementary to the *osbH* gene and the downstream region and by western blot using an anti-GFP-antibody.

To create OSBP7 and OSBP8 GFP-overexpressing cells, *osbG* and *osbH* were amplified from cDNA using the primers oMIB20 (*osbG* forward 5’CGA GAT CTA AAA TGG AGG CCG ATC CG), oMIB18 (*osbG* reverse with stop 5’ CCA CTA GTT TAA TTA CTA CCA CTT GCA GC), oMIB19 (*osbG* reverse without stop 5’ CCA CTA GTA TTA CTA CCA CTT GCA GC), oMIB21 (*osbH* forward 5’ CGA GAT CTA AAA TGT TTT CAG GAG CAT TG), oMIB23 (*osbH* reverse with stop 5’ CCA CTA GTT TAA TTT GAA GCT GCT GC) and oMIB22 (*osbH* reverse without stop 5’ CCA CTA GTA TTT GAA GCT GCT GCT TTA AC) and cloned into pDM317 and pDM323 (33) to generate N-and C-terminal GFP-fusions, respectively.

All plasmids used in this study are listed in Table S1. Plasmids were electroporated into *D. discoideum* and selected with the appropriate antibiotic. Hygromycin was used at a concentration of 50 μg/ml, blasticidin at a concentration of 5 μg/ml, and neomycin at a concentration of 5 μg/ml.

### SDS-PAGE and western blot

5×10^5^ cells were harvested and incubated with 2x Laemmli buffer containing ß-mercapto-ethanol and DTT. After the electrophoresis, proteins were transferred to a nitrocellulose membrane (Amersham^TM^ Protran^TM^, Premium 0,45µm NC) as described in (34). Transfer was performed for 50 min and a constant voltage of 120 V on a Mini Trans-Blot Cell (Biorad R) system. The membranes were stained with Ponceau S solution to check the efficiency of the protein transfer. For immunodetection, the membranes were blocked using non-fat dry milk and stained with an anti-GFP primary (Roche; 1:1000) and a goat anti-mouse secondary antibody coupled to horseradish peroxidase (HRP) (BioRad, 1:5000). The detection of HRP was accomplished using the Pierce^TM^ ECL Western Blotting Substrate (Thermo Scientific). The quantification of the band intensity was performed with ImageJ and GraphPad Prism.

### Cell viability assay

Cell viability was assessed by measuring the fraction of propidium iodide^+^ cells by flow cytometry. To this end, approximately 10^6^ cells were harvested and resuspended in Soerensen buffer (SB). Membrane damage was induced by addition of 5 mM LLOMe and measured in SB buffer containing 3 µM PI (Thermo Fisher Scientific). After one hour of incubation, 10,000 cells per condition were analysed using a SonySH800 and the PE-A channel. Flow cytometry plots were generated with FloJo.

### Induced pluripotent stem cell-derived macrophages (iPSDMs) differentiation and cell culture

iPSDM were generated from human induced pluripotent stem cell line KOLF2 sourced from Public Health England Culture Collections as previously described (13). To collect the cells, iPSDM were washed in 1x PBS and incubated with Versene (Gibco) for 10 min at 37 °C and 5 % CO_2_. Versene was diluted 1:3 in 1x PBS and cells were gently scraped, centrifuged at 300 *g*, resuspended in X-Vivo 15 (Lonza) supplemented with 2 mM Glutamax (Gibco), 50 μM β-mercaptoethanol (Gibco) and plated for experiments on 96-well CellCarrier^TM^ Ultra glass-bottom plates (Perkin Elmer) at approximately 50,000 cells per well.

### Mycobacteria strains, culture and plasmids

All the *M. marinum* material is listed in Table S1. *M. marinum* was cultured in 7H9 supplemented with 10 % OADC, 0.2 % glycerol and 0.05 % Tween-80 at 32 °C at 150 rpm until OD_600_ of 1 (∼1.5×10^8^ bacteria/ml). To prevent bacteria from clumping, flasks containing 5 mm glass beads were used. Luminescent *M. marinum* wt as well as ΔRD1 and ΔCE bacteria expressing mCherry were generated in the Thierry Soldati laboratory (27, 35, 36) and grown in medium supplemented with 25 µg/ml kanamycin and 100 µg/ml hygromycin, respectively. To generate wt and ΔRD1 mycobacteria expressing eBFP, the unlabelled strains were transformed with the pTEC18 plasmid (addgene #30177) and grown in medium with 100 µg/ml hygromycin.

All the *M. tuberculosis* material is listed in Table S1. *M. tuberculosis* were thawed and cultured in Middle 7H9 supplemented with 0.05% Tween-80, 0.2% glycerol and 10% ADC.

### Infection assays

The infection of *D. discoideum* with *M. marinum* was carried out as previously described (16, 37). Briefly, for a final MOI of 10, 5 x 10^8^ bacteria were washed twice and resuspended in 500 μl Hl5c filtered. To remove clumps, bacteria were passed 10 times through a 25-gauge needle and added to a 10 cm petri dish of *D. discoideum* cells. To increase the phagocytosis efficiency, the plates were centrifuged for two times 10 min at RT. After 20-30 min incubation, the extracellular bacteria were removed by several washes with Hl5c filtered. Finally, the infected cells were taken up in 30 ml of Hl5c at a density of 1 x 10^6^ c/ml supplemented with 5 µg/ml streptomycin and 5 U/ml penicillin to points, samples were taken for downstream experiments.

Infection of iPSDM was performed as previously described (13). Briefly, *M. tuberculosis* was grown to OD_600_ ∼ 0.8 and centrifuged at 2000 g for 5 min. The pellet was washed twice with PBS, shaken with 2.5-3.5 mm glass beads for 1 min to produce a single-bacteria suspension. Bacteria were resuspended in 8 ml of cell culture media and centrifuged at 300 g for 5 min to remove clumps. Bacteria were diluted to an MOI of 2 for infection before adding to the cells. After 2 hrs, the inoculum was removed, cells were washed with PBS, and fresh medium was added.

### Intracellular growth assays

*M. marinum* growth was assessed with the help of bacteria expressing luciferase as well as its substrates as previously described (35). Briefly, infected *D. discoideum* cells were plated in dilutions between 0.5 – 2.0 x 10^5^ on non-treated 96-well plates (X50 LumiNunc, Nunc) and covered with a gas permeable moisture barrier seal (4Ti). Luminescence was measured at 25 °C every hour for around 70 hrs using an Infinite 200 pro M-plex plate reader (Tecan).

### RNA-sequencing and proteomic data

RNA-Seq (15) and proteomics data (14) from *M. marinum* infected *D. discoideum* were re-analysed for selected genes involved in ER-contact site formation or lipid transport. All the data can be accessed via the supplementary files on BioRxiv.

RNA sequencing data of *M. tuberculosis*-infected macrophages was extracted from an original study (13). All RNA-Seq data is deposited in Gene Expression Omnibus (accession number GSE132283).

### Live cell imaging

To monitor non-infected cells or the course of infection by SD live imaging, cells were transferred to either 4- or 8-well μ-ibidi slides and imaged in low fluorescent medium (LoFlo, Formedium, UK) with a Zeiss Cell observer spinning disc (SD) microscope using the 63x oil objective (NA 1.46). To improve signal-to-noise, indicated images were deconvolved using Huygens Software from Scientific Volume Imaging (Netherlands). The images were further processed and analysed with ImageJ.

To analyse whether MCVs have impaired lysosomal or proteolytic function, infected cells were transferred to an ibidi slide and incubated for 10 min in Hl5c filtered medium containing 1µM LysoSensor Green (Thermo Fisher Scientific) or 1 hr with 50 µg/ml DQ-BSA Green (Thermo Fisher Scientific). In the case of LysoSensor Green labelling, the extracellular dye was removed before imaging. Z-stacks of 15 slices with 300 nm intervals were acquired.

To visualize GFP-Vps32, P4C-mCherry, P4C-GFP and OSBP8-GFP on damaged lysosomes, sterile membrane damage was induced with 5 mM LLOMe (Bachem) as described in (6). To label all endosomes, above mentioned cells were pre-incubated on ibidi slides overnight with 10 µg/ml dextran (Alexa Fluor™ 647, 10.000 MW,Thermo Fisher Scientific) in Hl5c filtered medium. Time acquired every 5 mins for at least 2 hrs.

For lattice light sheet microscopy (LLSM) the infection was performed as previously described. At 3 hours post infection (hpi) cells were seeded on 5 mm round glass coverslips (Thermo Scientific) and mounted on a sample holder specially designed for LLSM, which was an exact home-built clone of the original designed by the Betzig lab (38). The holder was inserted into the sample bath containing Hl5c filtered medium at RT. A three-channel image stack was acquired in sample scan mode through a fixed light sheet with a step size of 190 nm which is equivalent to a ∼189.597 nm slicing. A dithered square lattice pattern generated by multiple Bessel beams using an inner and outer numerical aperture of the excitation objective of 0.48 and 0.55, respectively, was used.

The raw data was further processed by using an open-source LLSM post-processing utility called LLSpy v0.4.9 (https://github.com/tlambert03/LLSpy) for deskewing, deconvolution, 3D stack rotation and rescaling. Deconvolution was performed by using experimental point spread functions and is based on the Richardson-Lucy algorithm using 10 iterations. Finally, image data were analysed and processed using imageJ and 3D surface rendering was performed with Imaris 9.5 (Bitplane, Switzerland).

### Antibodies, fluorescent probes, immunofluorescence and expansion microscopy

Fluoresbrite 641 nm Carboxylate Microspheres (1.75 µm) were obtained from Polysciences Inc., LysoSensor Green DND-189 as well as DQ Green BSA, Alexa Fluor 647 10kDa dextran and FM4- 64 from Thermo Fisher Scientific.

The anti-vatA, anti-vacA, anti-p80 antibodies were obtained from the Geneva antibody facility (Geneva, Switzerland). The anti-PDI antibody was provided from the Markus Maniak lab (University of Kassel, Germany). Anti-Ub (FK2) was from Enzo Life Sciences, the anti-OSBP antibody from Sigma-Aldrich. As secondary antibodies, goat anti-rabbit, anti-mouse and anti-rat IgG coupled to Alexa546 (Thermo Fisher Scientific), CF488R (Biotium), CF568 (Biotium) or CF640R (Biotium) were used.

For immunostaining of *D. discoideum*, cells were seeded on acid-cleaned poly-L-Lysine coated 10 mm coverslips and centrifuged at 500 g for 10 min at RT. Cells were fixed with 4 % paraformaldehyde/ picric acid and labelled with antibodies as described in (39). Images were acquired using an Olympus LSM FV3000 NLO microscope with a 60x oil objective with a NA of 1.40. Five slices with 500 nm intervals were taken.

Filipin staining was performed as previously described (18). Briefly, fixed cells were treated with Filipin at 50 µg/ml for 2 hrs without further permeabilization prior the primary antibody labelling. To avoid bleaching, images were taken using the SD microscope. Up to 20 slices with 300 nm intervals were obtained. All images were analysed and processed using ImageJ and graphical representations were generated using Graphpad Prism.

For immunostaining of infected iPSDMs, cells were fixed overnight with 4 % paraformaldehyde at 4 °C. Samples were quenched with 50 mM NH_4_Cl for 10 min and then permeabilized with 0.3 % Triton-X for 15 min. After blocking with 3% BSA for 30 min, samples were incubated with the anti-OSBP antibody for 1 hr at RT. After incubation, the coverslips were washed with PBS, before addition of the secondary antibody (45 min at RT). Nuclei were stained with DAPI. Images were recorded either with a Leica SP8 or an Opera Phenix (Perkin Elmer) with 63x water objective with a NA of 1.15.

The ExM protocol was adapted from (40) and (41). Briefly, cells were fixed with −20°C cold methanol and stained with antibodies as described before. The signal of mCherry and GFP was enhanced using a rat mAb anti-RFP (Chromotek, 5f8-100) and a rabbit pAb anti-GFP antibody (BIOZOL/MBL, MBL-598), respectively. Samples were then incubated with 1 mM methylacrylic acid-NHS (Sigma Aldrich) in PBS for 1 hr at RT in a 24-well plate. After washing three times with PBS, coverslips were incubated in the monomer solution (8.6% sodium acrylate, 2.5% acrylamide, 0.15% N,N’-methylenebisacrylamide, and 11.7% NaCl in PBS) for 45 min. This was followed by an 2 hrs incubation in the gelling solution (monomer solution, 4-hydroxy-TEMPO (0.01%), TEMED (0.2%) and ammonium persulfate (0.2%)) inside the humidified gelation chamber at 37 °C. Afterwards, gels were transferred into a 10-cm dish containing the digestion buffer (50 mM Tris, 1 mM EDTA, 0.5% Triton-X-100, 0.8M guanidine HCl, and 16 U/ml of proteinase K; pH 8.0) and incubated at 37 °C overnight. For final expansion of the polymer, gels were incubated in deionized water for at least 2.5 hrs. Subsequently, a region of interest was cut out and transferred onto a coverslip coated with poly-L-lysine to prevent movements during the imaging (Olympus LSM FV3000 NLO). Deionized water was used as imaging buffer and to store the samples at 4°C.

### CLEM with high-pressure freezing and freeze substitution

Cells expressing GFP-ABD and AmtA-mCherry or OSBP8-mCherry were seeded on poly-L-lysine sapphire discs (3 mm x 0.16 mm). Before seeding cells, a coordinate system was applied on the sapphires by gold sputtering using a coordinate template. Sapphires were dipped into 2% glutaraldehyde (GA) in HL5c and imaged in 0.5% GA in HL5c. Directly after acquisition of the LM image using the SD microscope, the sapphire discs were high-pressure frozen with a Compact 03 (M. Wohlwend, Switzerland) high pressure freezer (HPF). For HPF, the sapphire discs were placed with the cells and gold spacer facing onto flat 3 mm-aluminum planchettes (M. Wohlwend GmbH, Switzerland), which were beforehand dipped into hexadecene (Merck, Germany). The assemblies were thereafter placed into the HPF-holder and were immediately high pressure frozen. The vitrified samples were stored in liquid nitrogen until they were freeze substituted.

For freeze substitution (FS), the aluminum planchettes were opened in liquid nitrogen and separated from the sapphire discs. The sapphire discs were then immersed in substitution solution containing 1% osmium tetroxide (Electron Microscopy Sciences, Germany), 0.1% uranyl acetate and 5% H_2_O in anhydrous acetone (VWR, Germany) pre-cooled to −90 °C .The FS was performed in a Leica AFS2 (Leica, Germany) following the protocol of 27 hrs at −90 °C, 12 hrs at −60 °C, 12 hrs at −30 °C and 1 hr at 0 °C, washed 5 times with anhydrous acetone on ice, stepwise embedded in EPON 812 (Roth, Germany) mixed with acetone (30% EPON, 60% EPON, 100% EPON) and finally polymerized for 48 hrs at 60 °C. Ultrathin sections of 70 nm and semithin sections of 250 nm were sectioned with a Leica UC7 ultramicrotome (Leica, Germany) using a Histo diamond-and 35° Ultra diamond knife (Diatome, Switzerland). Sections were collected on formvar-coated copper slot grids and post-stained for 30 min with 2% uranyl acetate and 20 min in 3 % lead citrate and analyzed with a JEM 2100-Plus (JEOL, Germany) operating at 200 kV equipped with a 20 mega pixel CMOS XAROSA camera (EMSIS Germany).

For transmission electron microscopy (TEM) tomography 250 nm thick sections were labelled with 10 nm Protein-A gold fiducials on both sides prior to post-staining. Double tilt series were acquired using the TEMography software (JEOL, Germany) at a JEM 2100-Plus (JEOL, Germany) operating at 200 kV and equipped with a 20-megapixel CMOS XAROSA camera (EMSIS, Muenster, Germany). The nominal magnification was 12000x with a pixel size of 0.79 nm. Double tilt tomograms were reconstructed using the back-projection algorithm in IMOD (42).

### Serial block face (SBF) - scanning electron microscopy (SEM)

After SD microscopy in gridded ibidi 8-well chambers, cells were fixed in 2% GA in HL5c. Subsequently, samples were processed via adapted version of the NCMIR rOTO-post-fixation protocol (43) and embedded in hard Epon resin, ensuring pronounced contrast and electron dose resistance for consecutive imaging. All procedures were performed in the ibidi dish. In brief, after fixation, samples were post-fixed in 2% osmiumtetroxide (Electron Microscopy Sciences) and treated with 1.5 % (w/v) potassium ferrocyanide (Riedel de Haen) in Hl5c for 30 min. After washing in ultrapure water, cells were incubated in 1 % (w/v) thiocarbohydrazide (Riedel de Haen) in water for 20 min, followed by an additional 2 % osmication step for 1 hr at RT. Samples were then incubated in 1 % tannic acid in water for 30 min, washed and submerged in 1% aqueous uranyl acetate overnight at 4°C. Cells were then brought up to 50°C, washed and incubated in freshly prepared Walton`s lead aspartate (Pb(NO3)2 (Carl-Roth), L-Aspartate (Serva), KOH (Merck)) for 30 min at 60°C. Subsequently, cells were dehydrated through a graded ethanol (Carl-Roth) series (50 %, 70 % and 90 %) on ice for 7 min each, before rinsing in anhydrous ethanol twice for 7 min and twice in anhydrous acetone (Carl-Roth) for 10 min at RT. Afterwards, cells were infiltrated in an ascending Epon:acetone mixture (1:3, 1:1, 3:1) for 2 hrs each, before an additional incubation in hard mixture of 100% Epon 812 (Sigma). Final curation was carried out in hard Epon with 3% (w/w) Ketjen Black (TAAB) at 60°C for 48 hr. Once polymerized, the μ-Dish bottom was removed via toluene melting from the resin block leaving behind the embedded cells and finder grid imprint. ROIs were trimmed based on the coordinates (250*250*250 µm³) and the sample blocks were glued to aluminium rivets using two-component conductive silver epoxy adhesive and additionally coated in a 20 nm thick gold layer. The rivet containing the mounted resin block was then inserted into the 3View2XP (Gatan, USA) stage, fitted in a JSM-7200F (JEOL, Japan) FE-SEM, and precisely aligned parallel to the voltage, high vacuum mode of 10 Pa, utilizing a 30 nm condenser aperture and a positive stage bias of 400 V. Imaging parameters were set to 2 nm pixel size, 1.1 μs dwell time, in between ablation of 30 nm and an image size of 10240×10240 pixels. Overall, an approximate volume of 20×20×7 μm (a 210 slices) was acquired. Image acquisition was controlled via Gatan Digital Micrograph software (Version 3.32.2403.0). Further post processing, including alignment, filtering and segmentations were performed in Microscopy Image Browser (Version 2.7 (44)). Endoplasmic reticulum was traced and segmented manually throughout the entire dataset, whereas bacteria, nucleus and vacuole were annotated semi-automatically via morphological 3D watershed. Correlation of light microscopic and EM datasets was performed in AMIRA (Version 2021.1, Thermo Fisher) by rendering and overlaying both volumes, utilizing GFP and eBFP signal as natural landmarks within the electron micrographs.

## SUPPLEMENTAL MATERIAL

**FIG S1.**
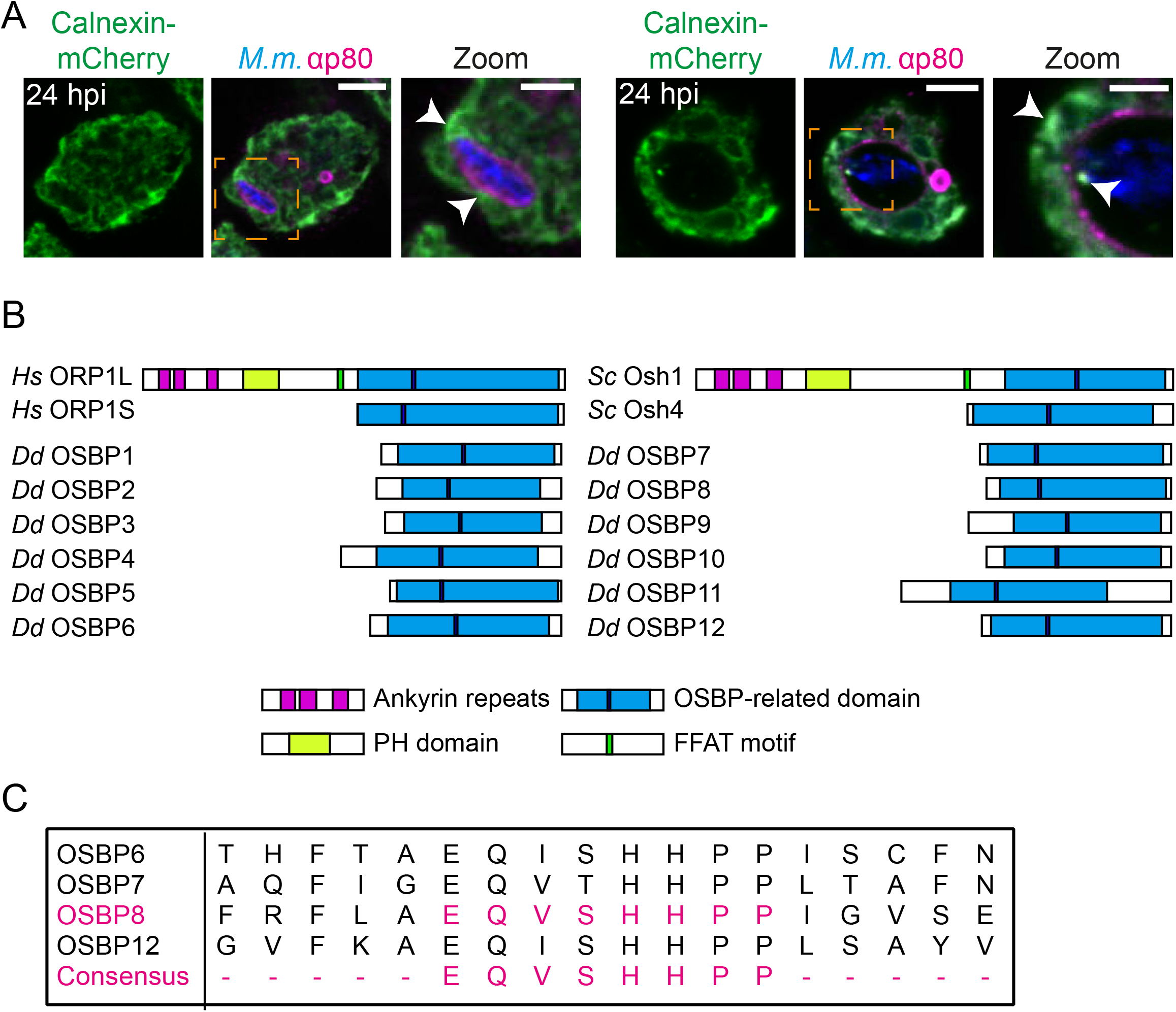
Mycobacterial infection induces ER-MCV contacts. (A) ER in apposition with the MCV. Representative images of cells showing calnexin-mCherry^+^ ER-tubules close to the MCV (arrow heads). Cells were infected with eBFP-expressing *M. marinum*. At 24 hpi cells were fixed and stained for p80 to label the membrane of the MCV. Scale bars, 5 µm; Zoom, 2 µm. (B) Domain organization of OSBPs in *D. discoideum* (*Dd*) compared to long (ORP1L, Osh1) and short ORP family members from *Homo sapiens* (*Hs*) and *Saccharomyces cerevisiae* (*Sc*) (ORP1S, Osh4). PH: pleckstrin homology; FFAT: two phenylalanines (FF) in an acidic tract. (C) Conserved fingerprint sequence of *D. discoideum* OSBPs.

**FIG S2.**
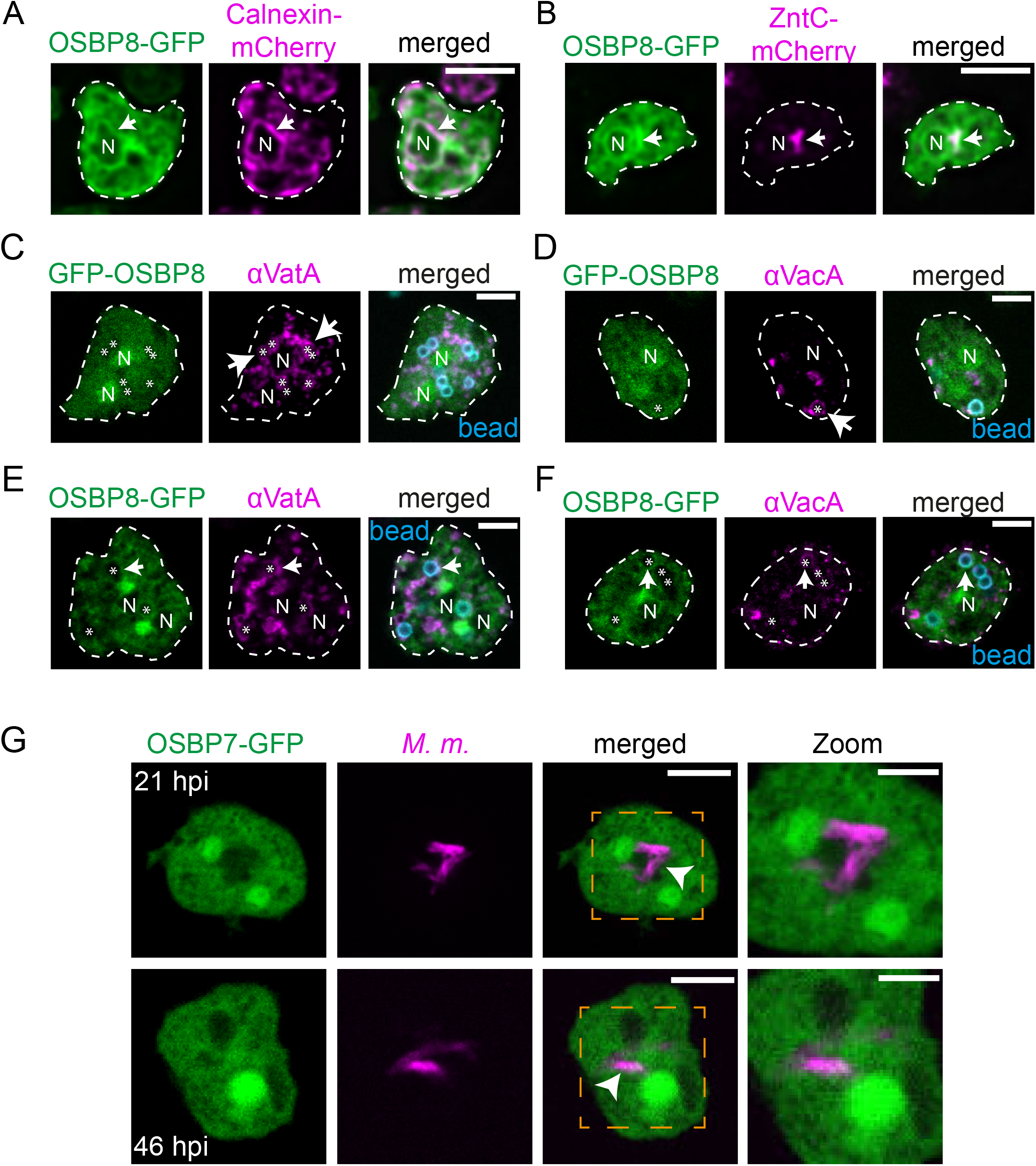
Localization of OSBP8-GFP and GFP-OSBP8 in non-infected cells. (A-B) OSBP8-GFP localizes in the cytosol, at the perinuclear ER and at the Golgi-apparatus. Cells overexpressing OSBP8-GFP/calnexin-mCherry or OSBP8-GFP/ZntC-mCherry were imaged live by SD microscopy. Arrow points to the juxtanuclear region or the Golgi-apparatus. Scale bars, 5 µm. Images were deconvolved. (C-D) GFP-OSBP8 locates in the cytosol and in the nucleus and is not mobilized to bead-containing phagosomes (BCPs). (E-F) OSBP8-GFP is not recruited to BCPs. Cells overexpressing OSBP8-GFP were incubated with fluorobeads for 2 hrs, fixed and then stained with αVatA (vATPase subunit A, lysosomes) and αVacA (VacuolinA, post-lysosomes) antibodies. Arrows point to VatA^+^or VacA^+^ BCPs. Asterisks indicate fluorobeads, N: nucleus. Scale bars, 5 µm. (G) OSBP7-GFP is not recruited to intracellular bacteria. Cells overexpressing OSBP7-GFP were infected with mCherry-expressing *M. marinum*. At the indicated time points, cells were imaged live by SD microscopy. Arrow heads indicate OSBP7-GFP^-^ mycobacteria. Scale bars, 5 µm; Zoom, 2 µm.

**FIG S3.**
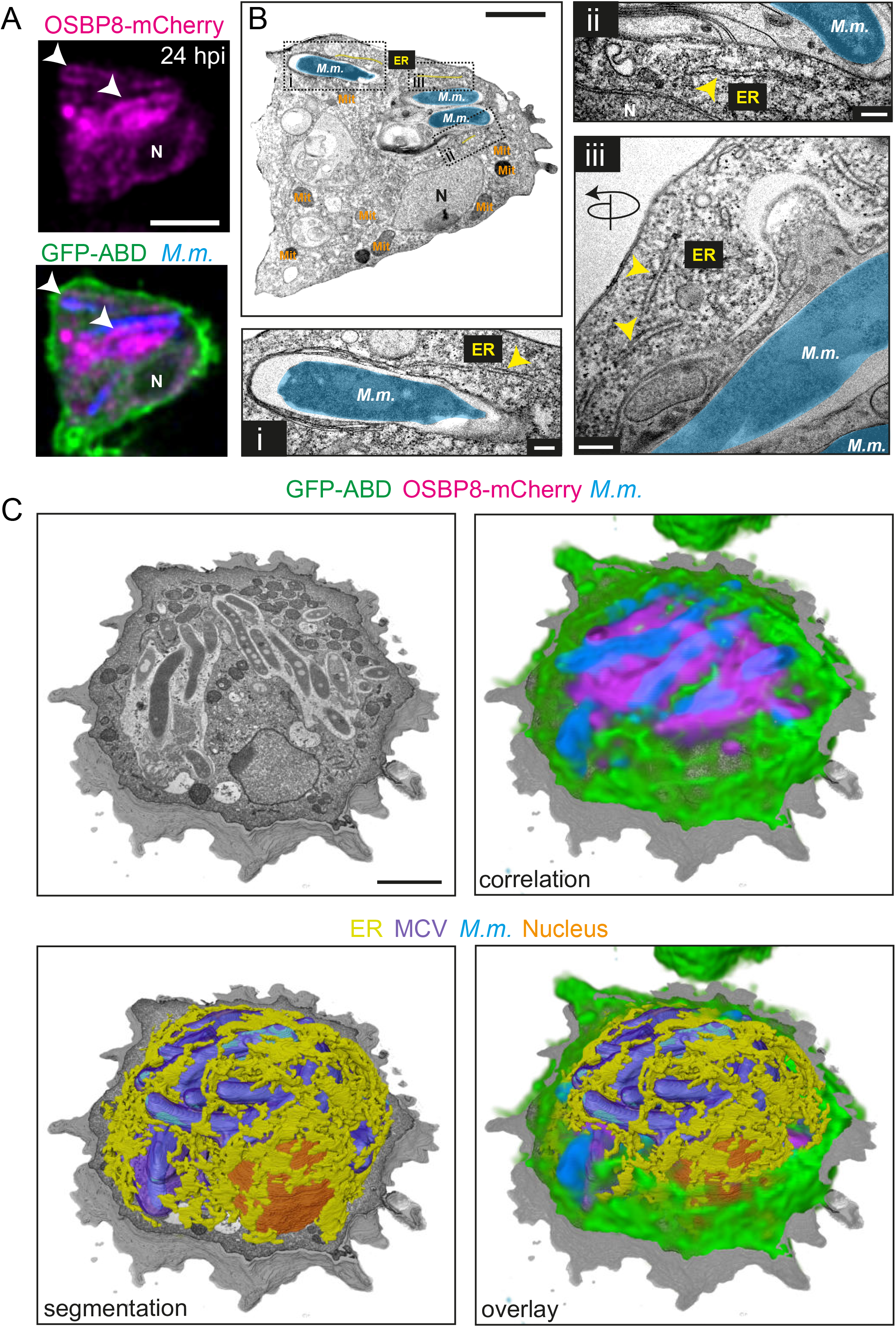
Correlative ultrastructural analysis revealed OSBP8-mCherry^+^ ER-tubules in close proximity to the MCV. (A) CLEM images showing OSBP8-mCherry at ER-MCV contacts (white arrow heads). Cells expressing OSBP8-mCherry/GFP-ABD were infected with eBFP-expressing *M. marinum*. At 24 hpi, cells were imaged after quick fixation by SD microscopy, high pressure frozen and prepared for EM. (B) EM micrograph. Positions of the closeups are indicated. (i - iii) Closeups showing OSBP8- mCherry^+^ ER-tubules close to the MCV. Yellow arrowheads point to ER-tubules in the vicinity of *M. marinum*. Mitochondria (Mit) were pseudo-coloured in orange, *M. marinum* (*M.m*.) in cyan and ER-tubules in yellow. N: nucleus. Scale bars, 5 µm (A); 2 µm (B) and 200 nm (i - iii). SD images were deconvolved. (C) SBF-SEM-derived images illustrate the correlation of the SD images and the volumetric segmentation of the EM data shown in Fig. 2F-G. The MCV is segmented in violet, *M. marinum* (*M.m*.) in cyan, ER-tubules in yellow and the nucleus in orange. Scale bars, 2 µm.

**FIG S4.**
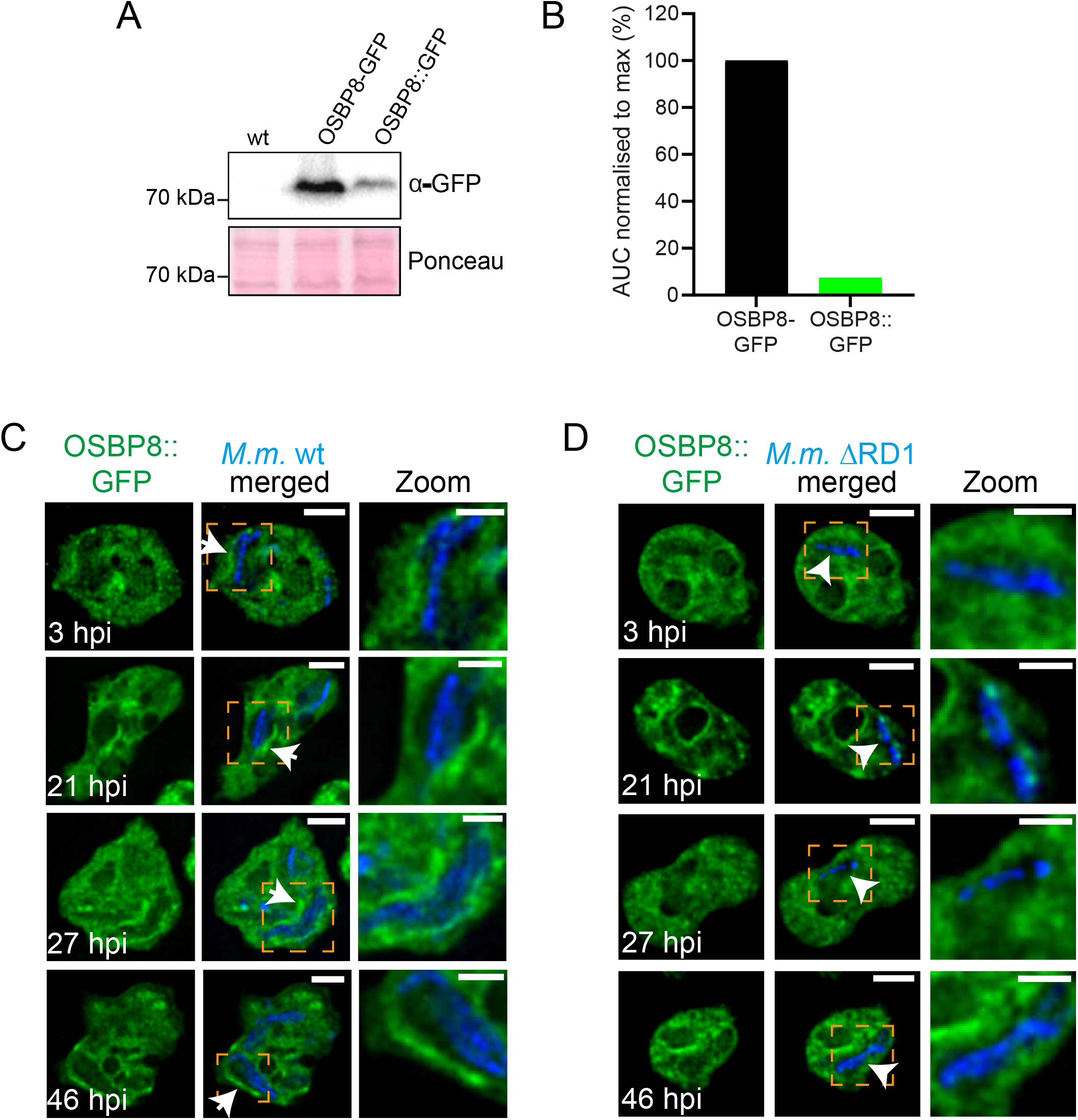
Endogenous OSBP8::GFP is mobilized during infection in an ESX-1 dependent manner. (A) Expression levels of OSBP8-GFP compared to OSBP8::GFP. (B) Quantification of A. Cells expressing OSBP8-GFP as well as OSBP8::GFP were harvested and then prepared for western blotting. The intensity of the bands was measured using ImageJ. AUC: area under the curve. (C-D) OSBP8::GFP mobilization during infection is dependent on ESX-1. Cells expressing OSBP8::GFP were infected with eBFP-expressing *M. marinum* wt or ΔRD1. At the indicated time points samples were taken for SD microscopy. Arrows point to OSBP8::GFP^+^ intracellular mycobacteria and arrow heads indicate OSBP8::GFP^-^ mycobacteria. Scale bars, 5 µm; Zoom, 2,5 µm. Images were deconvolved. *M.m.: M. marinum*.

**FIG S5.**
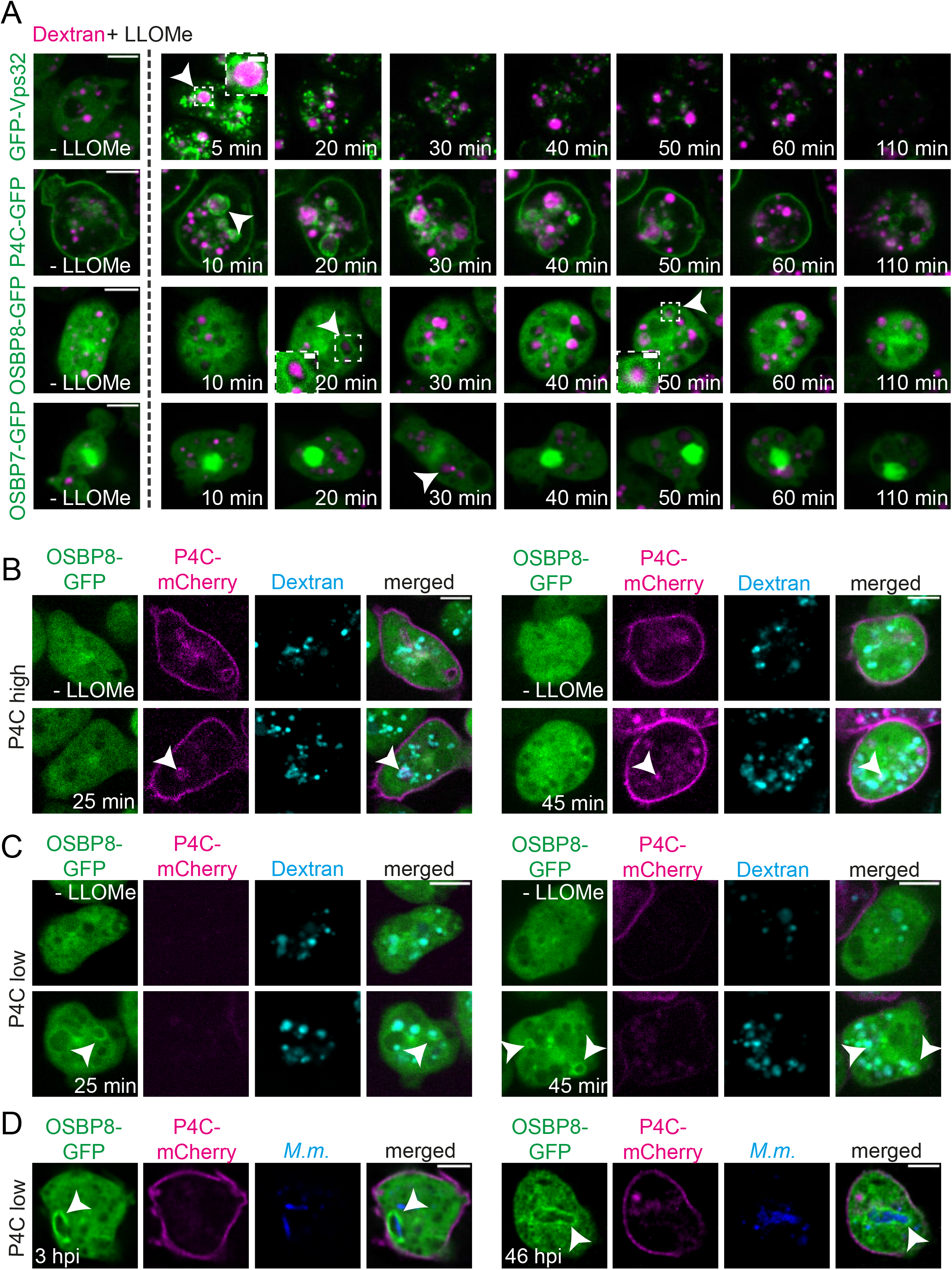
Dynamics of GFP-Vps32, P4C-GFP, OSBP8-GFP and OSBP7-GFP on damaged lysosomes. (A) Damaged lysosomes are positive for GFP-Vps32, P4C-GFP and OSBP8-GFP but not OSBP7-GFP. (B-C) In cells highly expressing P4C-mCherry, OSBP8-GFP is not recruited to damaged lysosomes and vice versa. Cells expressing GFP-Vps32, P4C-GFP, OSBP8-GFP, OSBP7-GFP and P4C-mCherry/OSBP8-GFP were incubated overnight with 10 kDa fluorescent dextran to label all endosomal compartments and then subjected to LLOMe. Arrow heads point to GFP-Vps32^+^, P4C-GFP^+,^ P4C-mCherry^+^, OSBP8-GFP^+^ or OSBP7-GFP^-^ lysosomes. Scale bars, 5 µm; Zoom, 1 µm. (D) OSBP8-GFP is recruited to MCVs of cells expressing P4C-mCherry at low levels. Cells overexpressing OSBP8-GFP/P4C-mCherry were infected with eBFP-expressing *M. marinum* wt. Images were taken at 3 and 46 hpi. Arrow heads point to OSBP8-GFP^+^ *M. marinum*. Scale bars, 5 µm. *M.m.: M. marinum*.

**FIG S6.**
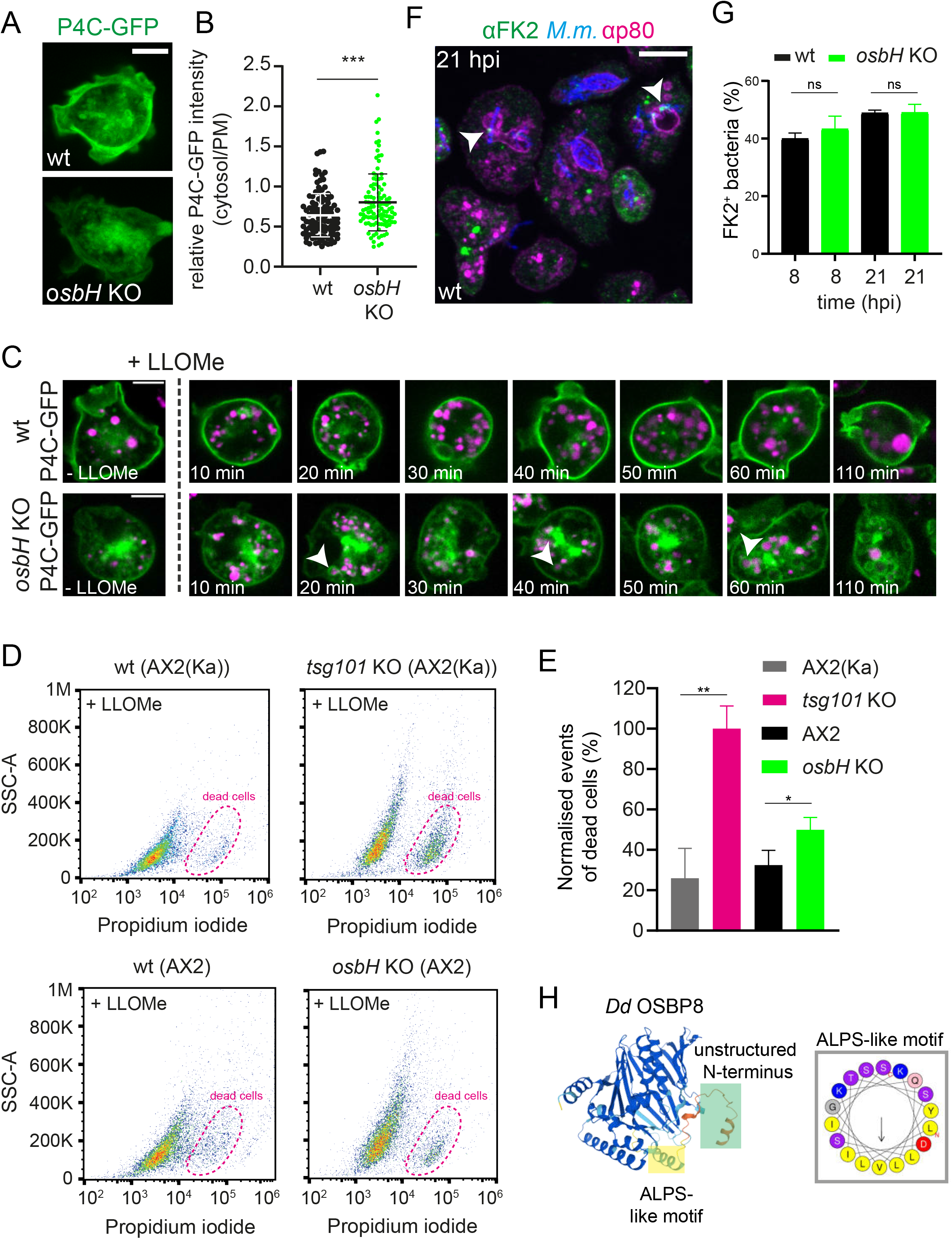
OSBP8 depletion leads to PI4P retention on damaged lysosomes and slightly affects cell viability but does not affect vacuolar escape of mycobacteria. (A) The PI4P distribution is altered in non-infected cells lacking OSBP8. (B) Quantification of A. Wt and *osbH* KO cells were imaged live. Shown are maximum z-projections of 15 z-stacks 300 nm apart. Scale bar, 5 µm. To label the PM for the quantification in B, cells were pre-stained with FM4-64. For each condition 108 cells per cell line were quantified using mageJ. The statistical significance of three independent experiments was calculated with an unpaired t-test (*** p<0.001). (C) P4C-GFP is not retrieved from damaged lysosomes in cells lacking OSBP8. Wt and *osbH* KO cells expressing P4C-GFP were incubated overnight with 10 kDa fluorescent dextran to label all endosomal compartments and then treated with LLOMe. GFP-signal of the *osbH* KO cells was enhanced for better visualisation. Scale bars, 5 µm. (D) Cell viability is affected in cells lacking OSBP8 upon LLOMe treatment. Wt, *osbH* or *tsg101* KOs were labelled with propidium iodide (PI) and incubated with LLOMe for 60 min. 10,000 cells were analysed per condition. Graphs are representative for two independent experiments. (E) Quantification of D. Plots were gated as indicated in D, to reveal the number of dead cells. Plots in E show the mean and standard deviation of three independent experiments. Statistical differences were calculated with a paired t test (*, P < 0.05; **, P < 0.01). (F) Vacuolar escape is unaltered in cells lacking OSBP8. (G) Percentage of ubiquitin^+^ bacteria in wt and *osbH* KOs at 8 and 21 hpi. Wt and *osbH* KO were infected with mCherry-expressing *M. marinum*, fixed and stained against ubiquitin (FK2) (green) and p80 (magenta). Representative maximum projections of 5 z-stacks of 500 nm at 21 hpi is shown in E. White arrowheads label ubiquitinated bacteria. Scale bars, 10 μm; Plots in G show the mean and standard deviation of three independent experiments (FK2 8, 21 hpi N=3, 138≤n≤462). Statistical differences were calculated with a paired t test.ns: not significant. *M.m.: M. marinum.* (H) OSBP8 contains an unstructured N-terminus as well as an ALPS-like motif. The OSBP8 structure was derived from AlphaFold (https://alphafold.ebi.ac.uk/entry/Q54QP6) and analysed using HeliQuest (https://heliquest.ipmc. cnrs.fr/).

**Movie S1** and **Movie S2** LLSM revealed that OSBP8-GFP^+^ membranes are capping the MCV (AmtA^+^). For more information see Fig. 2A-C.

**Movie S3** SBF-SEM. For more information see Fig. 2E-G, Fig. S3C.

## Acknowledgments

We greatly acknowledge the integrated Bioimaging facility (iBiOs) at the University of Osnabrück and especially Rainer Kurre, Michael Holtmannspötter and Olympia Ekaterini Psathaki for their expertise and friendly support. We thank Joost Holthuis for inspiring this project; Xiaoli Ma for cloning the *osbH* knockout constructs; and Ana T. López Jiménez, Jason King as well as Thierry Soldati for carefully reading this manuscript and their thoughtful suggestions. This work was supported by the Deutsche Forschungsgemeinschaft (SFB944-P25 (CB), SFB944-Z, SFB1557-P1 (CB), SFB1557-Z). The Barisch lab is a member of the SPP2225. This work was also supported by the Francis Crick Institute (to MGG), which receives its core funding from Cancer Research UK (FC001092), the UK Medical Research Council (FC001092), and the Welcome Trust (FC001092). This project has received funding from the European Research Council (ERC) under the European Union’s Horizon 2020 research and innovation programme (MGG, grant agreement n° 772022). For the purpose of open access, the author has applied a CC BY public copyright licence to any Author Accepted Manuscript version arising from this submission.

## Conflict of interests

The authors declare that they have no conflict of interest.

**Table S1.** Material used in this publication.

| *D. discoideum* Strains | Relevant characteristics | Source/Reference |
| --- | --- | --- |
| Ax2 | wt, parental strain of *osbH* KO, *osbH* KI |  |
| Ax2 *osbH* KO | Bsr<sup>r</sup> | This study |
| Ax2(Ka) | wt, parental strain of the *tsg101* KO |  |
| Ax2(Ka) *tsg101* KO | Bsr<sup>r</sup> | (20) |
| *D. discoideum* Plasmids |  | Source/Reference |
| OSBP8-GFP | pDM323-*osbH*, G418<sup>r</sup>, Amp<sup>r</sup> | This study |
| GFP-OSBP8 | pDM317-*osbH*, G418<sup>r</sup>, Amp<sup>r</sup> | This study |
| Calnexin-mCherry | pDM1044-*Calnx*, Hyg<sup>r</sup>, Amp<sup>r</sup> | (22) |
| ZntD-mCherry | pDM1044-*zntD*, Hyg<sup>r</sup>, Amp<sup>r</sup> | (20) |
| AmtA-mCherry | pDM1044-*amtA*, Hyg<sup>r</sup>, Amp<sup>r</sup> | (18) |
| OSBP8-mCherry | pDM1210-*osbH*, Hyg<sup>r</sup>, Amp<sup>r</sup> | This study |
| OSBP7-GFP | pDM323-*osbG*, G418<sup>r</sup>, Amp<sup>r</sup> | This study |
| GFP-OSBP7 | pDM317-*osbG*, G418<sup>r</sup>, Amp<sup>r</sup> | This study |
| P4C-mCherry | pDM1044-*p4c*, Hyg<sup>r</sup>, Amp<sup>r</sup> | (19),(22) |
| P4C-GFP | pDM323-*p4c*, G418<sup>r</sup>, Amp<sup>r</sup> | (45) |
| GFP-Vps32 | pDM317-*vps32*, G418<sup>r</sup>, Amp<sup>r</sup> | (6) |
| OSBP8::GFP | pPI183-*osbH*, Hyg<sup>r</sup>, Amp<sup>r</sup> | This study |
| GFP-ABD | pDXA-GFP-ABD120, G418<sup>r</sup>, Amp<sup>r</sup> | (46) |
| Mammalian cells | Relevant characteristics | Source/Reference |
| Human induced pluripotent stem cell-derived macrophages (iPSDMs) | iPSDMs were generated from human induced pluripotent stem cell line KOLF2 | Public Health England Culture Collections (catalogue number 77650100) |
| *M. marinum* material |  |  |
| *M. marinum* M | wt, parental strain | L. Ramakrishnan (University of Cambridge) |
| *M. marinum* ΔRD1 | RD1 locus deletion mutant | L. Ramakrishnan (University of Cambridge) (47) |
| *M. marinum* ΔCE | *esxA* and *esxB* deletion mutant | T. Soldati (University of Geneva) (27) |
| *M. tuberculosis* material |  |  |
| *M. tuberculosis* | H37Rv | Douglas Young (The Francis Crick Institute, London, UK), (13) |
| *M. tuberculosis* ΔRD1 | H37Rv ΔRD1 | Suzie Hingley-Wilson (University of Surrey, Guilford, UK), (13) |
| Mycobacteria Plasmids |  |  |
| pTEC18 | eBFP2 under control of the MSP promoter, Hyg<sup>r</sup>, Amp<sup>r</sup> | Addgene #30177(48) |
| pCherry10 | mCherry under control of the G13 promoter, Hyg<sup>r</sup>, Amp<sup>r</sup> | Addgene #24664 (49) |
| pTEC19 | E2-Crimson under the control of the MSP promoter, Hyg<sup>r</sup>, Amp<sup>r</sup> | Addgene #30178 (48) |

## REFERENCES

1. Uribe-Querol E, Rosales C. 2017. Control of Phagocytosis by Microbial Pathogens. Front Immunol 8:1368.

2. Hanna N, Koliwer-Brandl H, Lefrançois LH, Kalinina V, Cardenal-Muñoz E, Appiah J, Leuba F, Gueho A, Hilbi H, Soldati T, Barisch C. 2021. Zn(2+) Intoxication of Mycobacterium marinum during Dictyostelium discoideum Infection Is Counteracted by Induction of the Pathogen Zn(2+) Exporter CtpC. mBio 12.

3. Stinear TP, Seemann T, Harrison PF, Jenkin GA, Davies JK, Johnson PD, Abdellah Z, Arrowsmith C, Chillingworth T, Churcher C, Clarke K, Cronin A, Davis P, Goodhead I, Holroyd N, Jagels K, Lord A, Moule S, Mungall K, Norbertczak H, Quail MA, Rabbinowitsch E, Walker D, White B, Whitehead S, Small PL, Brosch R, Ramakrishnan L, Fischbach MA, Parkhill J, Cole ST. 2008. Insights from the complete genome sequence of Mycobacterium marinum on the evolution of Mycobacterium tuberculosis. Genome Res 18:729–41.

4. Cardenal-Munoz E, Barisch C, Lefrancois LH, Lopez-Jimenez AT, Soldati T. 2017. When Dicty Met Myco, a (Not So) Romantic Story about One Amoeba and Its Intracellular Pathogen. Front Cell Infect Microbiol 7:529.

5. Foulon M, Listian SA, Soldati T, Barisch C. 2022. Chapter 6 - Conserved mechanisms drive host-lipid access, import, and utilization in Mycobacterium tuberculosis and M. marinum, p 133–161. *In* Fatima Z, Canaan S (ed), Biology of Mycobacterial Lipids doi:https://doi.org/10.1016/B978-0-323-91948-7.00011-7. Academic Press.

6. López-Jiménez AT, Cardenal-Muñoz E, Leuba F, Gerstenmaier L, Barisch C, Hagedorn M, King JS, Soldati T. 2018. The ESCRT and autophagy machineries cooperate to repair ESX-1-dependent damage at the Mycobacterium-containing vacuole but have opposite impact on containing the infection. PLoS Pathog 14:e1007501.

7. Raykov L, Mottet M, Nitschke J, Soldati T. 2022. A TRAF-like E3 ubiquitin ligase TrafE coordinates endolysosomal damage response and cell-autonomous immunity to *Mycobacterium marinum*. bioRxiv doi:10.1101/2021.06.29.450281:2021.06.29.450281.

8. Barisch C, Holthuis JCM, Cosentino K. 2023. Membrane damage and repair: a thin line between life and death. Biological Chemistry doi:doi:10.1515/hsz-2022-0321.

9. Ellison CJ, Kukulski W, Boyle KB, Munro S, Randow F. 2020. Transbilayer Movement of Sphingomyelin Precedes Catastrophic Breakage of Enterobacteria-Containing Vacuoles. Curr Biol 30:2974–2983.e6.

10. Niekamp P, Scharte F, Sokoya T, Vittadello L, Kim Y, Deng Y, Südhoff E, Hilderink A, Imlau M, Clarke CJ, Hensel M, Burd CG, Holthuis JCM. 2022. Ca(2+)-activated sphingomyelin scrambling and turnover mediate ESCRT-independent lysosomal repair. Nat Commun 13:1875.

11. Radulovic M, Wenzel EM, Gilani S, Holland LK, Lystad AH, Phuyal S, Olkkonen VM, Brech A, Jäättelä M, Maeda K, Raiborg C, Stenmark H. 2022. Cholesterol transfer via endoplasmic reticulum contacts mediates lysosome damage repair. Embo j 41:e112677.

12. Tan JX, Finkel T. 2022. A phosphoinositide signalling pathway mediates rapid lysosomal repair. Nature 609:815–821.

13. Bernard EM, Fearns A, Bussi C, Santucci P, Peddie CJ, Lai RJ, Collinson LM, Gutierrez MG. 2020. M. tuberculosis infection of human iPSC-derived macrophages reveals complex membrane dynamics during xenophagy evasion. J Cell Sci 134.

14. Guého A, Bosmani C, Soldati T. 2019. Proteomic characterization of the *Mycobacterium marinum*- containing vacuole in *Dictyostelium discoideum*. bioRxiv doi:10.1101/592717:592717.

15. Hanna N, Burdet F, Melotti A, Bosmani C, Kicka S, Hilbi H, Cosson P, Pagni M, Soldati T. 2019. Time-resolved RNA-seq profiling of the infection of *Dictyostelium discoideum* by *Mycobacterium marinum* reveals an integrated host response to damage and stress. bioRxiv doi:10.1101/590810:590810.

16. Hagedorn M, Soldati T. 2007. Flotillin and RacH modulate the intracellular immunity of Dictyostelium to Mycobacterium marinum infection. Cell Microbiol 9:2716–33.

17. Tailleux L, Neyrolles O, Honoré-Bouakline Sp, Perret E, Sanchez Fo, Abastado J-P, Lagrange PH, Gluckman JC, Rosenzwajg M, Herrmann J-L. 2003. Constrained Intracellular Survival of Mycobacterium tuberculosis in Human Dendritic Cells 1. The Journal of Immunology 170:1939–1948.

18. Barisch C, Paschke P, Hagedorn M, Maniak M, Soldati T. 2015. Lipid droplet dynamics at early stages of Mycobacterium marinum infection in Dictyostelium. Cell Microbiol 17:1332–49.

19. Vormittag S, Hüsler D, Haneburger I, Kroniger T, Anand A, Prantl M, Barisch C, Maaß S, Becher D, Letourneur F, Hilbi H. 2023. Legionella-and host-driven lipid flux at LCV-ER membrane contact sites promotes vacuole remodeling. EMBO Rep doi:10.15252/embr.202256007:e56007.

20. Barisch C, Kalinina V, Lefrançois LH, Appiah J, López-Jiménez AT, Soldati T. 2018. Localization of all four ZnT zinc transporters in Dictyostelium and impact of ZntA and ZntB knockout on bacteria killing. J Cell Sci 131.

21. Repnik U, Borg Distefano M, Speth MT, Ng MYW, Progida C, Hoflack B, Gruenberg J, Griffiths G. 2017. L-leucyl-L-leucine methyl ester does not release cysteine cathepsins to the cytosol but inactivates them in transiently permeabilized lysosomes. J Cell Sci 130:3124–3140.

22. Steiner B, Swart AL, Welin A, Weber S, Personnic N, Kaech A, Freyre C, Ziegler U, Klemm RW, Hilbi H. 2017. ER remodeling by the large GTPase atlastin promotes vacuolar growth of Legionella pneumophila. EMBO Rep 18:1817–1836.

23. Weber SS, Ragaz C, Reus K, Nyfeler Y, Hilbi H. 2006. Legionella pneumophila exploits PI(4)P to anchor secreted effector proteins to the replicative vacuole. PLoS Pathog 2:e46.

24. Ragaz C, Pietsch H, Urwyler S, Tiaden A, Weber SS, Hilbi H. 2008. The Legionella pneumophila phosphatidylinositol-4 phosphate-binding type IV substrate SidC recruits endoplasmic reticulum vesicles to a replication-permissive vacuole. Cell Microbiol 10:2416–33.

25. Mesmin B, Bigay J, Polidori J, Jamecna D, Lacas-Gervais S, Antonny B. 2017. Sterol transfer, PI4P consumption, and control of membrane lipid order by endogenous OSBP. Embo j 36:3156-3174.

26. Fineran P, Lloyd-Evans E, Lack NA, Platt N, Davis LC, Morgan AJ, Hoglinger D, Tatituri RV, Clark S, Williams IM, Tynan P, Al Eisa N, Nazarova E, Williams A, Galione A, Ory DS, Besra GS, Russell DG, Brenner MB, Sim E, Platt FM. 2016. Pathogenic mycobacteria achieve cellular persistence by inhibiting the Niemann-Pick Type C disease cellular pathway. Wellcome Open Res 1:18.

27. Cardenal-Munoz E, Arafah S, Lopez-Jimenez AT, Kicka S, Falaise A, Bach F, Schaad O, King JS, Hagedorn M, Soldati T. 2017. Mycobacterium marinum antagonistically induces an autophagic response while repressing the autophagic flux in a TORC1- and ESX-1-dependent manner. PLoS Pathog 13:e1006344.

28. Nakatsu F, Kawasaki A. 2021. Functions of Oxysterol-Binding Proteins at Membrane Contact Sites and Their Control by Phosphoinositide Metabolism. Front Cell Dev Biol 9:664788.

29. Chung J, Torta F, Masai K, Lucast L, Czapla H, Tanner LB, Narayanaswamy P, Wenk MR, Nakatsu F, De Camilli P. 2015. INTRACELLULAR TRANSPORT. PI4P/phosphatidylserine countertransport at ORP5- and ORP8- mediated ER-plasma membrane contacts. Science 349:428-32.

30. Drin G, Casella JF, Gautier R, Boehmer T, Schwartz TU, Antonny B. 2007. A general amphipathic alpha-helical motif for sensing membrane curvature. Nat Struct Mol Biol 14:138–46.

31. Faix J, Kreppel L, Shaulsky G, Schleicher M, Kimmel AR. 2004. A rapid and efficient method to generate multiple gene disruptions in Dictyostelium discoideum using a single selectable marker and the Cre-loxP system. Nucleic Acids Res 32:e143.

32. Paschke P, Knecht DA, Williams TD, Thomason PA, Insall RH, Chubb JR, Kay RR, Veltman DM. 2019. Genetic Engineering of Dictyostelium discoideum Cells Based on Selection and Growth on Bacteria. J Vis Exp doi:10.3791/58981.

33. Veltman DM, Akar G, Bosgraaf L, Van Haastert PJ. 2009. A new set of small, extrachromosomal expression vectors for Dictyostelium discoideum. Plasmid 61:110–8.

34. Towbin H, Staehelin T, Gordon J. 1979. Electrophoretic transfer of proteins from polyacrylamide gels to nitrocellulose sheets: procedure and some applications. Proc Natl Acad Sci U S A 76:4350–4.

35. Arafah S, Kicka S, Trofimov V, Hagedorn M, Andreu N, Wiles S, Robertson B, Soldati T. 2013. Setting up and monitoring an infection of Dictyostelium discoideum with mycobacteria, p 403-17. *In* Eichinger L, Rivero F (ed), Dictyostelium Protocols (Methods Mol Bio), vol 983. Humana Press.

36. Kicka S, Trofimov V, Harrison C, Ouertatani-Sakouhi H, McKinney J, Scapozza L, Hilbi H, Cosson P, Soldati T. 2014. Establishment and validation of whole-cell based fluorescence assays to identify anti-mycobacterial compounds using the Acanthamoeba castellanii-Mycobacterium marinum host-pathogen system. PLoS One 9:e87834.

37. Barisch C, Lopez-Jimenez AT, Soldati T. 2015. Live Imaging of Mycobacterium marinum Infection in Dictyostelium discoideum. Methods Mol Biol 1285:369–85.

38. Chen BC, Legant WR, Wang K, Shao L, Milkie DE, Davidson MW, Janetopoulos C, Wu XS, Hammer JA, 3rd, Liu Z, English BP, Mimori-Kiyosue Y, Romero DP, Ritter AT, Lippincott-Schwartz J, Fritz-Laylin L, Mullins RD, Mitchell DM, Bembenek JN, Reymann AC, Böhme R, Grill SW, Wang JT, Seydoux G, Tulu US, Kiehart DP, Betzig E. 2014. Lattice light-sheet microscopy: imaging molecules to embryos at high spatiotemporal resolution. Science 346:1257998.

39. Hagedorn M, Neuhaus EM, Soldati T. 2006. Optimized fixation and immunofluorescence staining methods for Dictyostelium cells. Methods Mol Biol 346:327–38.

40. Chen F, Tillberg PW, Boyden ES. 2015. Optical imaging. Expansion microscopy. Science 347:543–8.

41. Chozinski TJ, Halpern AR, Okawa H, Kim HJ, Tremel GJ, Wong RO, Vaughan JC. 2016. Expansion microscopy with conventional antibodies and fluorescent proteins. Nat Methods 13:485–8.

42. Kremer JR, Mastronarde DN, McIntosh JR. 1996. Computer visualization of three-dimensional image data using IMOD. J Struct Biol 116:71–6.

43. Deerinck T, Bushong E, Lev-Ram V, Shu X, Tsien R, Ellisman M. 2010. Enhancing Serial Block-Face Scanning Electron Microscopy to Enable High Resolution 3-D Nanohistology of Cells and Tissues. Microscopy and Microanalysis 16:1138–1139.

44. Belevich I, Joensuu M, Kumar D, Vihinen H, Jokitalo E. 2016. Microscopy Image Browser: A Platform for Segmentation and Analysis of Multidimensional Datasets. PLOS Biology 14:e1002340.

45. Welin A, Weber S, Hilbi H. 2018. Quantitative Imaging Flow Cytometry of Legionella-Infected Dictyostelium Amoebae Reveals the Impact of Retrograde Trafficking on Pathogen Vacuole Composition. Appl Environ Microbiol 84.

46. Pang KM, Lee E, Knecht DA. 1998. Use of a fusion protein between GFP and an actin-binding domain to visualize transient filamentous-actin structures. Curr Biol 8:405–8.

47. Volkman HE, Clay H, Beery D, Chang JC, Sherman DR, Ramakrishnan L. 2004. Tuberculous granuloma formation is enhanced by a mycobacterium virulence determinant. PLoS Biol 2:e367.

48. Takaki K, Davis JM, Winglee K, Ramakrishnan L. 2013. Evaluation of the pathogenesis and treatment of Mycobacterium marinum infection in zebrafish. Nat Protoc 8:1114–24.

49. Carroll P, Schreuder LJ, Muwanguzi-Karugaba J, Wiles S, Robertson BD, Ripoll J, Ward TH, Bancroft GJ, Schaible UE, Parish T. 2010. Sensitive detection of gene expression in mycobacteria under replicating and non-replicating conditions using optimized far-red reporters. PLoS One 5:e9823.

